# Glycans in fertilization and adhesion: histochemical and ultrastructural insights from the chaetognath *Spadella cephaloptera*

**DOI:** 10.1101/2025.10.09.681355

**Authors:** Cristian Camilo Barrera Grijalba, Sabine Thetter-Dürr, Julian Bibermair, Tim Wollesen

**Author notes:** To whom correspondence should be addressed: Tim Wollesen, Cristian Camilo Barrera Grijalba.

## Abstract

**Background:** To cope with strong and rapidly changing water currents, some marine invertebrates have evolved complex adhesive mechanisms that involve the interplay of different biomolecules, such as carbohydrates. Carbohydrates may, however, also be involved in other physiological processes such as reproduction, a research field poorly studied in protostomes. The benthic chaetognath *Spadella cephaloptera* is a protandric hermaphrodite capable of rapid attachment and detachment to substrates. Nevertheless, the putative underlying glycobiology during the adhesion process but also other physiological processes remain unknown for chaetognaths.

**Methods:** In the present study, through histological, immunohistochemical, and ultrastructural analysis, we characterized the location of different glycans in the reproductive and attachment systems of *S. cephaloptera.* Developmental changes of adhesive cells were investigated from early hatchlings up to adults.

**Results:** Acidic and sulfated mucosubstances were detected in the sperm ducts, whereas mature oocytes were surrounded by a carboxylated jelly coat. The distribution of adhesive cells in *S. cephaloptera* shifted from anterior discrete individual cells in hatchlings to posteriorly distributed cell clusters in the adult. Moreover, we identified secretion granules enriched in fibrous content inside the adhesive cells. Furthermore, immunohistochemistry revealed microtubule processes linking the adhesive cells to the intraepidermal plexus. Lectin affinity tests showed strong apical PNA binding and clear PHA-L/E and ConA signals in the adhesive cells. No evidence for a dual gland detachment system was observed.

**Conclusions:** This study provides the first glycan distribution analysis in a chaetognath, revealing the presence of carbohydrates in different structures of the reproductive system, highlighting their putative relevance during the fertilization process. Furthermore, the observed glycan moieties suggest that *S. cephaloptera* also combines convergently evolved features reported for other temporary attachment systems in marine invertebrates. However, *S. cephaloptera* also displays unique characters, such as specific ontogenetic changes occurring during early developmental stages that correspond to its feeding behavior. Our observations do not support the presence of additional gland cells mediating detachment. Finally, this work sets the framework for a molecular characterization of the reproductive and attachment systems of the enigmatic chaetognaths.

## Background

Intertidal organisms are exposed to steep, rapidly fluctuating gradients of salinity, temperature, pH, and strong water currents [1]. Research on the distribution of biomolecules such as carbohydrates in the cells of marine invertebrates has provided insights into the unique mechanisms they have evolved, which underpin their ecological success in rapidly changing habitats [2, 3, 4]. Chaetognaths (arrow worms) are marine predators essential for the maintenance of a variety of larger forms of life [5, 6, 7]. The phylogenetic position of this phylum within Metazoa remains contested as they exhibit a mosaic of characters observed in divergent taxa [8, 9]. In this view, chaetognaths have been suggested to be part of protostomes due to the organization of the germ layers during early development and features of the nervous system, such as the presence of a circumesophageal brain [5, 10, 11]. Nevertheless, they also feature the formation of the mouth opposite to the blastopore during gastrulation, a tripartite body architecture, and the co-expression of the gene (*brachyury*) around the blastopore and the stomodeum, suggesting deuterostome affinity [5, 12]. In this regard, research on chaetognaths requires data on fundamental cell processes, such as those mediated by carbohydrates concerning reproduction and attachment, as this information could provide insight into evolutionary comparisons.

The best-studied chaetognath to date is *Spadella cephaloptera*, a marine benthic predator. It exhibits a temporary attachment system, claimed to be mediated by clusters of ventrally elongated cells in the posterior region of the body [13]. In addition, these cells display electron-dense secretory granules in the apical region when examined using a transmission electron microscope (TEM) [5]. Temporary attachment mechanisms have been described in a variety of marine invertebrates, including representatives from echinoderms, mollusks, chordates, and cnidarians [14, 15, 16]. In these organisms, the presence of lectin-detectable glycans has been demonstrated, as evidenced by Peanut agglutinin (PNA) labeling of the Galβ1-3GalNAc motifs present in the attachment cells of *Macrostomum lignano*, *Ciona intestinalis*, and *Asterina gibbosa* [17, 18, 19]. Notably, it has been described that subsets of these glycans can be part of secreted proteins, establishing specific types of glycosylation that confer electrostatic properties to the glue [2]. No comparable histochemical data exist for *S. cephaloptera*, nor has it been described how the attachment process may change during the ontogeny of this chaetognath. Considering the above, the analysis of the glycan architecture of the adhesive system of *S. cephaloptera*, besides providing insight into its underlying mechanism, may also offer a venue for extending the taxonomic scope of bio-adhesion and provide information on potentially conserved mechanisms related to wet adhesion.

Research on the bio-adhesion of marine invertebrates has shown that secretions from these organisms have the potential to generate solutions in various fields, including hydrogel formation, wet glues, and insights for biofouling prevention [15, 20]. Extensive research has been done on the content of the byssus thread of bivalves, demonstrating the presence of the modified amino acid 3,4-dihydroxyphenyl-l-alanine (DOPA) in the proteinaceous fraction [21]. This amino acid has been utilized as the core of multiple bio-inspired applications, providing a route for designing materials that mimic its chemical properties in the search for a product with the original properties of bivalve glue [22]. However, it has also been demonstrated that while DOPA properties are relevant for the attachment, some other players have been understudied [23, 24]. Furthermore, research on other marine invertebrates is not as advanced, and many taxa potentially featuring relevant solutions are still not being investigated. Therefore, the research conducted on the attachment system of *S. cephaloptera* provides the opportunity to uncover additional properties of wet adhesives that are of interest in various branches of research.

Like all chaetognaths*, S. cephaloptera* is a protandric hermaphrodite and, as observed also in other members of Spadellidae, exhibits elaborate mating behaviors [25]. Former studies have mostly focused on fertilization and reproductive processes, relying primarily on ultrastructural data, but information regarding the molecules potentially regulating these processes is still missing [5, 26]. Moreover, evidence on the glycan composition of sexual gametes in chaetognaths is still missing. In addition, in the sea urchin these glycans have been demonstrated to play an essential role during fertilization, mediating gamete recognition and triggering downstream mechanisms such as the release of cortical secretory granules to prevent polyspermy [27, 28]. Furthermore, in *Parasagitta elegans*, it is known that fluctuations in whole-body lipid and carbohydrate levels overlap with their reproductive cycles [29], yet the role of polysaccharides within the reproductive system of chaetognaths remains unknown. Therefore, histological characterization of the polysaccharides present in the reproductive cells may provide insight into how these molecules could be related to the reproduction of *S. cephaloptera*.

Notably, it has been observed that changes in environmental conditions, such as pH, affect the fertilization success and capabilities of marine invertebrates, including sea urchins [30]. In this case, the dynamics of successful interactions between the oocytes and sperm have been demonstrated to be compromised in acidic conditions [30]. Given the role of glycans in the fertilization process in other marine invertebrates and the essential role that chaetognaths play in the aquatic food webs, the understanding of the potential role of carbohydrates in the reproduction of *S. cephaloptera* raises interest, considering the drastic changes in pH and temperature levels the sea environment is undergoing [31].

In this study, we examined the reproductive and attachment systems of *S. cephaloptera* by histochemical, immune histochemistry and ultrastructural analyses. Our results suggest a potential sperm-supporting role of sulfated and carboxylated polysaccharides in the seminal fluid contained in the ducts of the testis in adults of *S. cephaloptera*. In addition, we observed an enrichment of exclusively carboxylated polysaccharides in mature oocytes, consistent with a role for carbohydrate-mediated gamete recognition. Regarding the adhesive system, our results support the idea that *S. cephaloptera* relies on an epithelial secretion to attach to the substrates. Specifically, vesicle-like vacuoles enriched in neutral glycans were found to be over-represented in the ventral papillae. We also observed accumulation of secretion granules in the apical region of the adhesive cells, which contained a fibrillar agent. Furthermore, the distribution of adhesive cells exhibits a stage-specific pattern matching the physiology of the chaetognath. The adhesive papillae are individuated cells during early stages and form distinct clusters of cells during adulthood. The distribution of acetylated α-tubulin, together with observations of synaptic vesicles in the ultrastructure of the adhesive cells, indicate a relationship between the attachment system and the intraepidermal nervous plexus. The latter is innervated from the ventral nerve center (VNC), hinting at the neurosecretory role of the adhesive cells. The ultrastructural data do not support that the detachment mechanism of *S. cephaloptera* involves specific secretory cells, as no features indicative of a dual gland system were observed. Lastly, by reviewing the distribution of lectin-binding glycans, we show that peanut agglutinin (PNA) exhibits high specificity for the apical region of the adhesive cells, suggesting that a Galβ1-3GalNAc glycoconjugate could be involved in the attachment mechanism as observed in other marine invertebrates with temporary attachment systems [18, 19, 32].

## Methods

### Animal collection and husbandry

Adult specimens of *Spadella cephaloptera* were collected at low tides using plankton nets on the shore adjacent to the Station Biologique de Roscoff (Roscoff, France), at the approximate location: 48°43’47.2"N, 3°59’15.3"W, in the summer of 2023. The specimens were placed in petri dishes with natural seawater at 35 ppt to induce mating and egg laying. Egg batches were transferred to separate petri dishes to record individual hatching times, and seawater was exchanged daily. Before fixation, all the studied specimens from the different stages of *S. cephaloptera* were relaxed in a solution containing 100 mM MgCl2 in filtered seawater and incubated at 4 °C, depending on their developmental stage, as follows: 24 hours post-hatching (hph) and 5 days post-hatching (dph) for 10 minutes; late juveniles and adults for 2 hours.

### Transmission electron microscopy

Specimens at 24 hph, 5 dph, and adults of *S. cephaloptera* were prepared according to established protocols with modifications [11]. The samples were fixed in Karnovsky’s solution for 2 hours in a water bath at 48 °C (3% glutaraldehyde, 1% paraformaldehyde, and 30% filtered seawater, containing 0.2M sodium cacodylate buffer pH 7.3. Afterward, the specimens were rinsed every 30 minutes for 2 hours using the cacodylate solution with an addition of 9% glucose. Post-fixation was done in 1% osmium tetroxide in distilled water for 1h at room temperature, followed by repeated rinsing and finally dehydration in an ascending ethanol series and 100% acetone. Later, the samples were gradually infiltrated with epoxy resin, Agar 100 (Agar Scientific) overnight on an orbital shaker at room temperature, using acetone as intermedium, embedded and polymerized at 65 °C for 70h. Subsequently, ribbons of 1 µm serial sections were produced using a Histo Jumbo diamond knife (Diatome) on a Leica Ultramicrotome UC6 (Leica Microsystems), as previously reported [33]. The generated sections were stained with a solution of 1% Toluidine Blue for 25 seconds at 80 °C. Stained sections were analyzed using a light microscope, and sections containing a region of interest, were resectioned to 60-80 nm following established protocols [34]. Ultra-thin sections were contrasted using 1% uranyl acetate for 8 minutes, followed by three rinses. Next, the grids were counterstained for 3 minutes in 3% lead citrate and rinsed thrice again. Ultrastructure of the ultra-thin sections was documented using a ZEISS Libra 120 transmission electron microscope (Zeiss). The resulting images were later processed using Fiji v. 2.16.0 [35] and IrfanView v. 4.72 [36].

### 3D reconstruction

The semi-thin serial sections of 24 hph and 5 dph specimens of *S. cephaloptera* were used for 3D reconstruction of the ventral epidermal papillae. The sections were imaged using a Nikon DS-Ri2 camera mounted on a Nikon Ni-U compound microscope (Nikon), The image stack was loaded into Fiji v.2.16.0 [35], where it was transformed into 8-bit greyscale images. Images unsuitable for reconstruction were deleted and replaced by adjoining images, using the stack sorter tool of FIJI. The resulting stack was registered in Amira v. 2022 (Thermo Scientific) and aligned using the AlingSlices module which manual adjustments. Adhesive papillae identified in the epidermal surface of *S. cephaloptera* were manually segmented in the Segmentation Editor. The surface of the papillae was generated following the steps of previous studies [37]. The reconstructed cells were visualized in the specimen via a volume rendering of the latter combined with a surface viewer module. Snapshots were exported for further image processing.

### Histological staining procedures

Adult specimens of *S. cephaloptera* were fixed for one hour in Bouin’s solution (72% picric acid, 4% formaldehyde, 24% glacial acetic acid) at room temperature. Later, the samples were washed four times for 15 minutes each with 70% ethanol and stored at 4 °C before further processing. The samples were dehydrated in an ascending ethanol series. Next, the specimens were transferred into benzol for 1 hour and finally incubated in paraffin at 60 °C overnight. Subsequently, paraffin blocks were stored at 4 °C for 16 hours before sectioning. Ribbons of thin sections of 6 μm were produced using the Leica RM2235 microtome (Leica) and later stained with the following staining procedures to identify the general properties of the glycans present in the tissue: Alcian Blue (AB) (pH = 2.5) for detecting carboxylated and sulfated polysaccharides; AB (pH = 1.0) for targeting exclusively sulfated polysaccharides, both with hematoxylin counterstain, and Periodic Acid Shiff reaction (PAS), for detecting neutral glycans [38, 39, 40]. Stained slides were sealed with DMX and documented using a Nikon DS-Ri2 camera mounted on a Nikon NI-U microscope.

### Immunohistochemistry

Hatchlings, juveniles, and adults were fixed overnight at 4 °C in a fixing solution (4% paraformaldehyde, 0.1 M MOPS, 2 mM MgSO4, 1 mM EGTA, 0.5 M NaCl). The samples were washed three times for 5 minutes with PBS-T (1X PBS, pH 7.4, and 0.01% Tween 20) and then stored in PBS with 0.05% Sodium Azide. Afterward, the specimens were stepped in PBTr (1x PBS pH 7.4 with 0.3% Triton X-100(v/v)) and washed three times for 10 minutes. From this point onwards, and up to sample mounting, incubations were performed at 4 °C. Unspecific antibody binding was treated using a blocking solution (PBTr supplemented with 3% (v/v) Normal Goat Serum, NGS; (Invitrogen) on a rotatory shaker at 70 rpm overnight. Subsequently, the samples were incubated in the primary antibody (anti-acetylated a-tubulin from mouse) (Sigma) diluted 1:2000 in the blocking solution overnight. The unbound primary antibody was washed three times with PBTr for 10, 30, and 60 minutes. Later, the samples were incubated overnight in a secondary antibody (anti-mouse Alexa 633) (Thermo Fischer Scientific) at 1:500 in PBTr supplemented with 1% NGS. The excess antibody was washed out using thrice PBTr for 60 minutes. At this point, the samples were gradually transferred to PBS-T (1x PBS, pH 7.4, with 0.05% Tween 20) and washed three times with the same solution for 10 minutes. Subsequently, to stain F-actin filaments, the samples were incubated overnight in PBS-T supplemented with Alexa Fluor® 488 phalloidin (Invitrogen) at a dilution of 1:40 and (0.3%) DAPI in the dark. The excess phalloidin and DAPI (Carl Roth) were removed using PBS-T washes, repeated 8 times for 10 minutes each. Next, the specimens were mounted on glass slides with Vectashield (Vector Laboratories) and visualized using a Leica Stellaris 5/DMi8 confocal microscope (Leica) with sequential fluorescence scans.

### Lectin affinity assay

Hatchlings (24 hph) of *S. cephaloptera* were fixed as described for the immunohistochemistry assays. Lectin fluorescence histochemistry was performed as previously described with modifications [19]. Briefly, the specimens were washed four times for 15 minutes in TBS-T (TRIS buffer, pH 8.0, supplemented with 5 mM CaCl2 and 0.1% Triton X-100). From this step onwards, all incubations are carried out at 4 °C. Unspecific binding was treated using a blocking solution (TBS-T with 3% (w/v) bovine serum albumin (Sigma-Aldrich) overnight. Subsequently, biotinylated lectins (Vector Laboratories) were added at a final concentration of 25 mg/ml and incubated overnight. Later, unbound lectins were washed using TBS-T six times for 10 min. To fluorescently label the attached lectins, the samples were incubated for 1 hour in the blocking solution with a 1:300 dilution of streptavidin conjugated to Alexa Fluor 633 (Sigma-Aldrich). The samples were washed eight times for 15 minutes with TBS-T and then incubated for 1 hour with DAPI at a ratio of 1:350 in TBS-T. After removing the excess stain with three washes of TBST-T for 5 minutes each, the samples were mounted in Vectashield and visualized using a Leica Stellaris 5 / DMi8 - confocal microscope (Leica).

## Results

### Carbohydrate distribution in the adult chaetognath *Spadella cephaloptera*

The distribution of acidic or carboxylated polysaccharides in adults of *Spadella cephaloptera* detected by AB (pH = 2.5), is found in distinctive reproductive structures but not in the adhesive cells (Fig.1). By reviewing the clusters of ventral adhesive cells in the tail region and in the posterior end of the trunk, it was not possible to identify staining derived from the binding of AB at pH 2.5 in the bodies of the adhesive cells or surrounding epidermal cells (Fig. 1A, B). In contrast, the sperm ducts located between the longitudinal muscles of the tail, feature the presence of acidic or carboxylated polysaccharides (Fig. 1A, C). In addition, by examining the posterior end of the tail, no staining was observed in the seminal vesicles (Fig. 1A, C). Moreover, it was observed that the distribution of acidic or carboxylated polysaccharides extends up to the posterior septum, which separates the trunk from the anterior part of the tail, involving the entire span of the duct (Fig. 1D).

**Figure 1.**
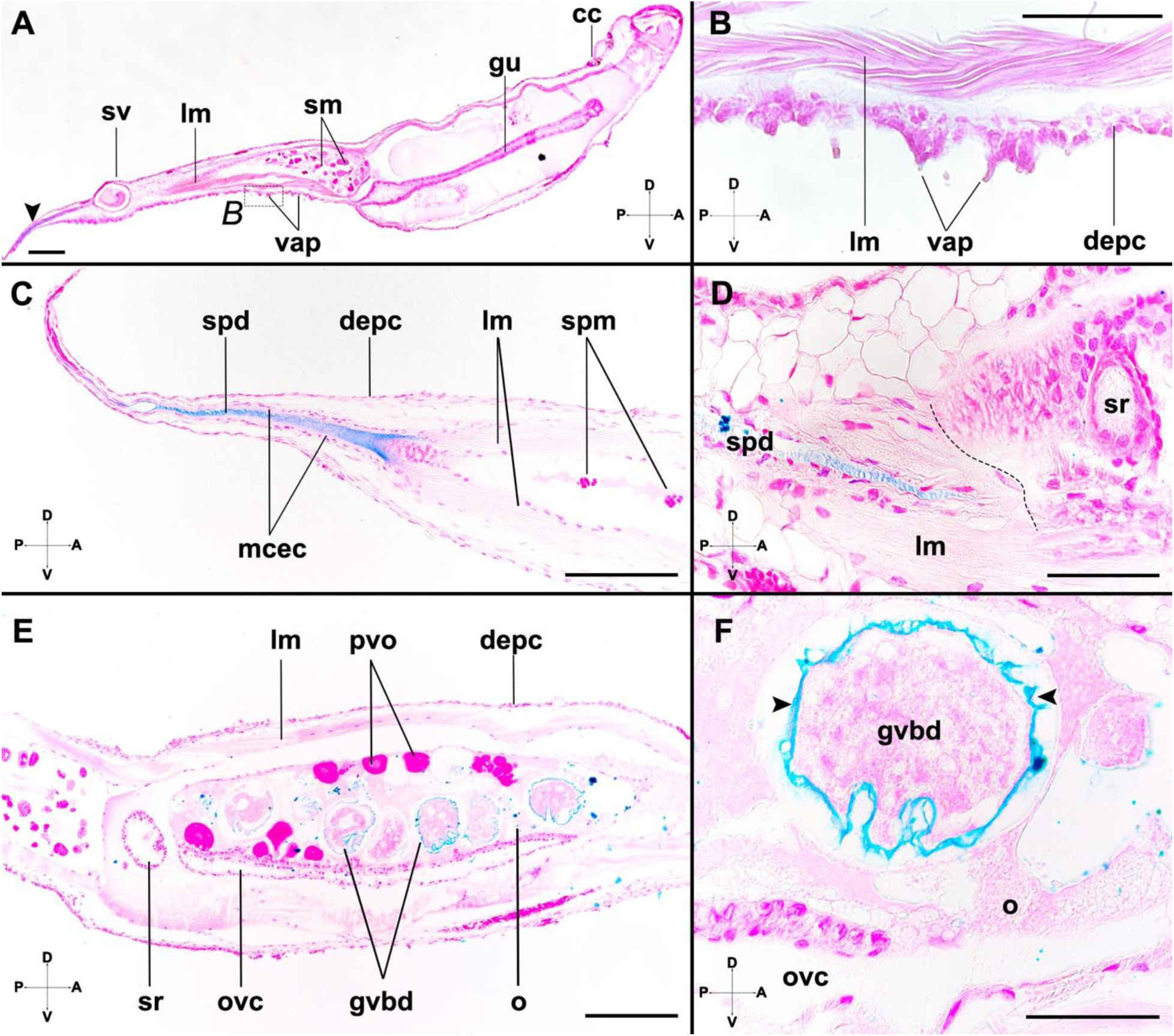
Location of carboxylated and sulfated polysaccharides in the structures of one adult specimen of *Spadella cephaloptera*. The cell nuclei are counterstained with hematoxylin (magenta) and the polysaccharides are stained with Alcian blue pH 2.5 (blue). **A**. Sagittal section displaying the overview of the coelomic cavities and internal organs and structures of *S. cephaloptera*. Stained polysaccharides are visualized in the posterior end of the tail (arrowhead), near the seminal vesicles (sv). **B.** Detailed aspect visualizing the ventral adhesive papillae (vap) that are surrounded by distal epidermal cells (depc) but devoid of evident staining related to the presence of detectable polysaccharides at pH 2.5. **C.** Close-up in the posterior end of the tail, where the sperm duct is enriched in sulfated and carboxylated polysaccharides. **D.** Limit between the tail coelom and the trunk cavities, displaying staining of Alcian blue up to the posterior septum (dotted line). **E.** Overview of the distribution of sulfated and carboxylated polysaccharides in the trunk cavities, which contains mature oocytes after the germinal vesicle breakdown (gvbd). Pre-vitellogenic oocytes (pvo) are absent in this region. **F.** Detail of one mature oocyte contained in the ovary exhibiting carboxylated and sulfated polysaccharides in the presumptive jelly coat (arrowheads). Additional abbreviations: gu, gut; corona ciliata, cc; longitudinal muscles, lm; mcec, multiciliated epithelial cells; o, ovary; ovc, oviducal complex; sm, spermatogonial masses; sr, seminal receptacles. Scale bars: A = 200 µm; B, D, F = 50 µm; C, E = 100 µm.

In the trunk region, the mature oocytes are located within the coelomic cavities [26]. They are surrounded by an AB^+^ (pH = 2.5) layer (Fig. 1E, F). Indeed, the staining forms a distinct coat that surrounds the membrane of the ovulated eggs that move later to the oviducal complex, which is part of the trunk cavities (Fig. 1F).

The location of the exclusively sulfated polysaccharides, as detected by AB (pH = 1.5), overlaps with the staining patterns observed at pH 2.5 (cf. Figs. 1-2). In this regard, the staining in the tail coelom is visible in the sperm ducts at a lower intensity; however, it was not observed inside the sperm vesicles (Fig. 2A-B). In contrast, in the trunk region, there was no evidence of acidic polysaccharides covering the mature oocytes (Fig. 2C). Staining was, however, observed inside the seminal receptacles, which are located bilaterally above the posterior septum (Fig. 2C-D). In the head region, the chitinous grasping spines that insert at the lateral plates exhibited AB^+^ (pH = 1.5) staining (Fig. 2E). Furthermore, the ventral adhesive papillae showed no detectable sulfated mucopolysaccharides, which suggests that the molecules inside the secretion granules may display different glycan moieties (Fig. 2F; cf. [5]).

**Figure 2.**
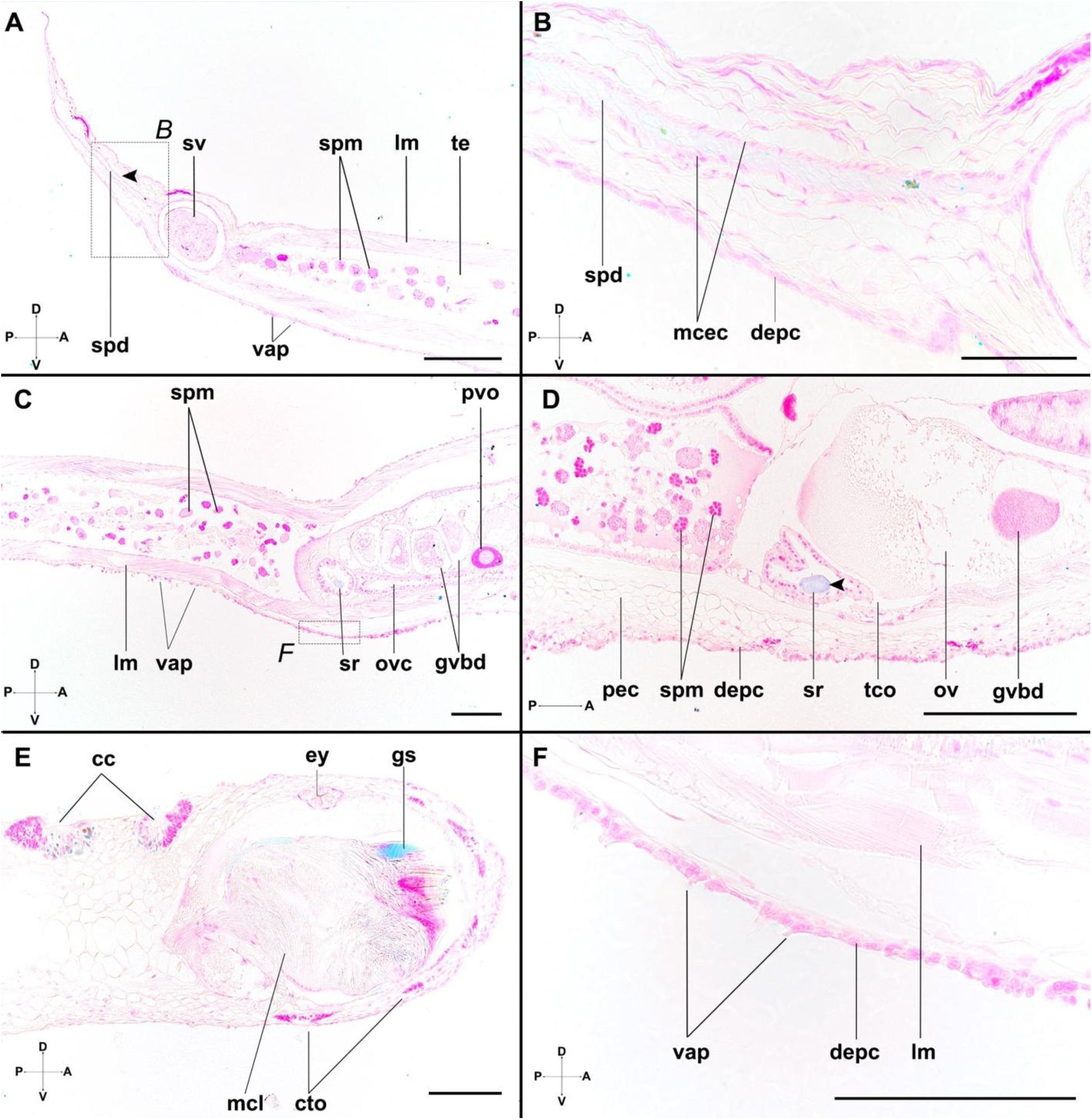
Distribution of sulfated polysaccharides in organs and structures in one adult specimen of *Spadella cephaloptera*. The cell nuclei are stained with hematoxylin (pink/magenta), and the glycans are visualized with Alcian blue pH 1 (blue/ cyan). **A.** Sagittal section of the posterior end of the tail, in which the sperm duct (spd) exhibits the presence of sulfated polysaccharides. **B.** Detail of the sperm duct (referenced in A), which is surrounded by multiciliated epithelial cells (mcec). **C.** Overview of the transition between trunk and tail, where there is staining located in the seminal receptacle (sr), and there is no evidence of sulfated polysaccharides in the jelly coat of oocytes after the germinal vesicle breakdown (gvbd), nor in the pre-vitellogenic oocytes (pvo). **D.** Frontal section showing a close-up of the seminal receptacle, which contains a defined vesicle with staining for sulfated glycans (arrowhead). **E.** Sagittal section of the anterior region of the head of the adult chaetognath, where the grasping spines around the vestibular musculature exhibit chitinous structures that are detected by Alcian blue pH 1. Notably, there is also staining derived from the ciliary structures of the corona ciliata (cc). **F.** Close-up of the ventral adhesive papillae (vap) from the tail (referenced in C), for which there is no evidence of the presence of acidic mucopolysaccharides. Additional abbreviations: cto, ciliary tuft organs; depc, distal epidermal cells; ey, eye; lm, longitudinal muscle; mcl, lateral muscle; ovc, oviducal complex; pec, proximal epidermal cells; spm, spermatogonial masses; sv, seminal vesicle; tco, trunk coelom; te, testis. Scale bars: A = 200 µm; B, F = 50 µm; C, D-E = 100 µm.

The neutral carbohydrates detected by the PAS reaction were localized in different body regions of *S. cephaloptera* (Fig. 3). In the tail region, the staining was observed in the sperm duct and included the testis accommodated within the tail coelomic cavities (Fig. 3A). Strongly stained vesicle-like structures were concentrated along the anterior surface of the tail. The glycoconjugates identified in this region appeared to co-localize with the cell clusters forming the ventral adhesive papillae (Fig. 3B–C). These neutral carbohydrates within the distal epidermal layer showed a posteriorly oriented distribution, extending toward the transition zone between trunk and tail. Strong PAS+ signal was also observed in the dorsally located corona ciliata and in the grasping spines (Fig. 3D). Notably in the anterior region of the epidermis, there was no evidence of staining that could match in size, aggregation density, and location the ventral adhesive papillae (Fig. 3A, D). Further structures with visible staining were not considered to be positive for the PAS reaction as they were also labeled in the negative control samples (Suppl. Fig. 1).

**Figure 3.**
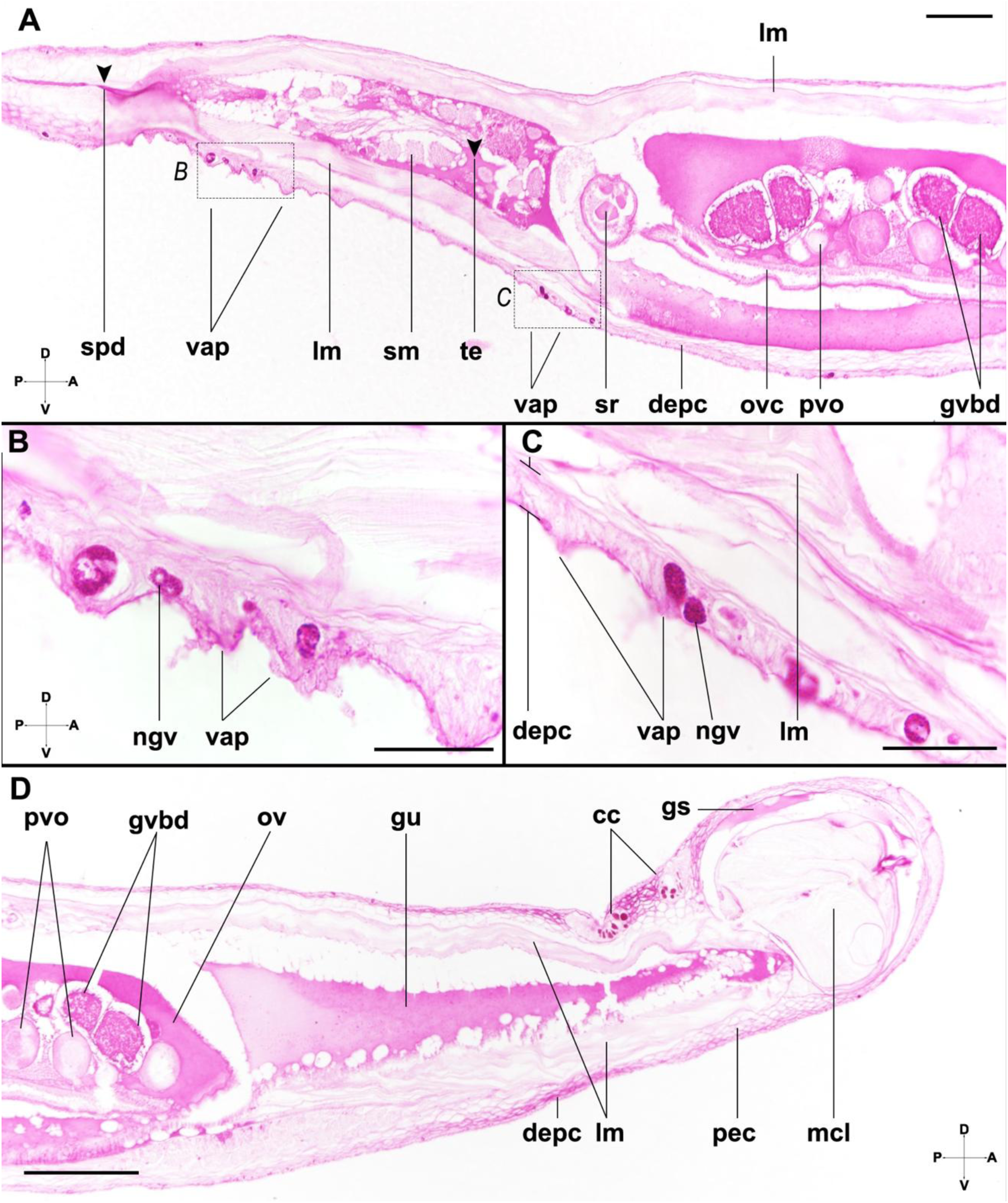
Organization of neutral carbohydrates detected by PAS staining in the adult of *Spadella cephaloptera* (magenta). **A.** Sagittal section of the mid-posterior region of the body highlighting staining (arrowheads) in the tail, the sperm duct (spd), and testis (te). **B.** Clusters of ventral adhesive papillae (vap) from the most outer layer of epidermis in the tail (referenced in A) containing strongly stained neutral glycan vesicles (ngv). **C.** Adhesive cells located below the posterior septum (noted in A), displaying dense vesicles (equivalent to the ones observed in **C.**). **D.** Anterior region of the body of *S. cephaloptera* in which the grasping spine exhibits carbohydrate staining and no groups of vesicles are detected in the distal (depc) or proximal epidermis regions (pec). Additional abbreviations: cc, corona ciliata; gs, grasping spine; gvbd, germinal vesicle break down; gu, gut; lm, longitudinal muscles; mcl, lateral muscle; ov, ovary; ovc, oviducal complex; pvo, previtellogenic oocyte; sm, spermatogonial masses; sr, seminal receptacles. Scale bars: A = 100 µm; B, C = 50 µm; D = 200 µm.

### Ultrastructure of the developing adhesive system

At 24 hph, specimens of *S. cephaloptera* exhibit ventral adhesive cell papillae with a pyramidal morphology, having approximately a base of 6,5 μm and a height of 9 μm up to the apical end (Fig. 4A). These cells are embedded in the distal region of the multilayered epidermis. In the proximal region of the adhesive cell, a nucleus occupies a significant portion of the cell with a transverse area of 95 μm^2^. In the cytoplasm, there is an abundance of filaments that are about the size of microtubules, ∼ 21nm (Fig. 4A, B) [41]. Numerous electron-dense granules are present in the body of the adhesive cells and are organized towards the apical region where they are supposedly secreted (Fig. 4A, B). Inside the adhesive papillae, secretory granules were observed near ribosomes, suggesting a possible role of these organelles in granule formation (Suppl. Fig. 2A). The surrounding epidermal cells envelop most of the cell body of the papilla, leaving an uncovered area of around 1.9 μm. This may point to a structural supportive role, which is mediated by these cells during attachment. Furthermore, at 24 hph, all adhesive cells are discretely distributed, with no evidence of physical contact between them (Fig. 4A).

**Figure 4.**
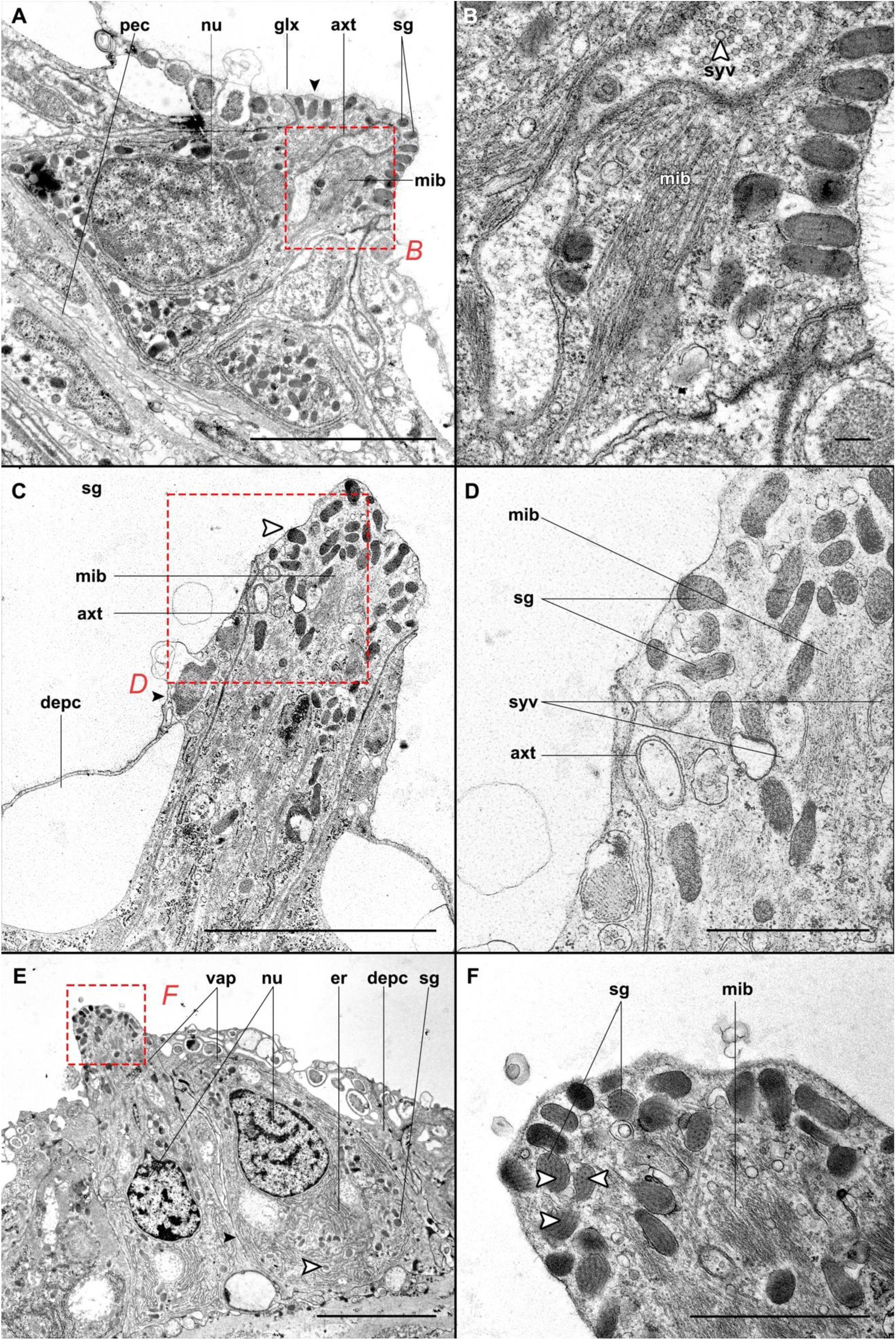
Transmission electron microscopy images of the ventral adhesive papillae during ontogeny. **A.** Overview of the ventral adhesive papillae at 24 hph, characterized by the pyramid elongated shape with a large nucleus in the basal region, surrounded by distal epidermal cells (depc) and exhibiting axonal terminals (axt) inside. **B**. Detail of the apical region of the adhesive cell (referenced in A), where secretion granules (sg) accumulate close to the membrane. Synaptic vesicles (syv) can be seen (arrowhead) within the axonal terminal. **C.** The ventral adhesive cell at 5dph exhibits similar features to the ones at 24 hph (A), but notably more elongated. The cell also contains electron-dense granules accumulated in the distal region (white arrowhead) and is surrounded by epidermal cells that are actively releasing material to the exterior (black arrowhead). **D.** Close-up of the adhesive cell apical region (referenced in C), where the microtubules (mib) and double-membrane axonal terminals are eventuated. **E.** Overview of two elongated adhesive cells in the adult of *S. cephaloptera* being in physical contact (arrowhead) and embedded in the distal epidermis region (depc). Note how the secretion granules are found in the basal region next to the endoplasmic reticulum (er) (white arrowheads). **F.** Close-up of the most apical part of the adhesive cell (noted in E), in which the fiber-like content inside the secretion granules is evident (arrowheads). Additional abbreviations. Scale bars: A, C, E = 5 µm; B = 200 nm; D, F = 1,25 µm.

At 5 dph, the size of each adhesive cell of *S. cephaloptera* increases about 1.3 times, forming a cell that rises approximately 12,2 μm from the most distal epidermis layer (Fig. 4C). The electron dense granules described above for 24 hph specimens, are similarly clustered close to the most apical part of the cell at 5 dph (Fig. 4C). In this regard, for 5 dph individuals, structures indicative of granule secretion were observed (Suppl. Fig. 2B), suggesting that their content may be actively secreted during ontogeny. A closer inspection of the secretory granules revealed parallel-oriented, strip-like structures, with a diameter of approximately 13 nm (Fig. 4C, D). No evidence of contact between the cell bodies of different adhesive papillae was found during this developmental stage.

When *S. cephaloptera* reaches adulthood, the most distal layer of the epidermis gets thicker (about 7 μm) and shows high secretory activity, resulting in the appearance of electron dense granules of multiple sizes in TEM (Suppl. Fig. 2C). The adhesive cells protrude around 16 μm from the base of the distal epidermis at this point in development. Of particular interest, this is the only stage in which we found evidence for the attachment cells forming clusters of at least two cells, and in which their membranes are in physical contact with each other (Fig. 4E). Additionally, the strip-like structures observed in the electron-dense secretory granules at 5dph can be also distinguished in adult samples, showing that this feature is conserved during ontogeny (Fig. 4F).

### Distribution of the adhesive papillae during ontogeny

The location of the adhesive cells in *Spadella cephaloptera* is stage-specific. At 24 hph, adhesive cells are present in the antero-ventral half of the body, starting from the most anterior part of the head in the cephalic rim (Fig. 5A-C). In this structure, there is an enrichment of attachment cells. In the forming neck region, there are few papillae which become more abundant towards the posterior region of the trunk (Fig. 5C). From a transversal section in the mid-trunk region, it becomes evident that adhesive papillae are not clustered in specific body regions of the transversal axis constituting a distinctive attachment organ but rather a multiple-point attachment system (Fig. 5D). In 5dph specimens, the spatial distribution of the attachment cells exhibited notable changes. In this sense, no evidence of an anterior cephalic rim or adhesive cells was found on the ventral surface of the head region (Fig. 5E-G). Instead, during this stage, the adhesive cells display an increased density in the trunk region, spreading posteriorly from the anterior beginning of the subjacent lateral somata clusters (Fig. 5E, H). Overall, in 5dph individuals, the adhesive cells are arranged in a discrete cell distribution in the trunk resembling the condition of 24 hph specimens.

**Figure 5.**
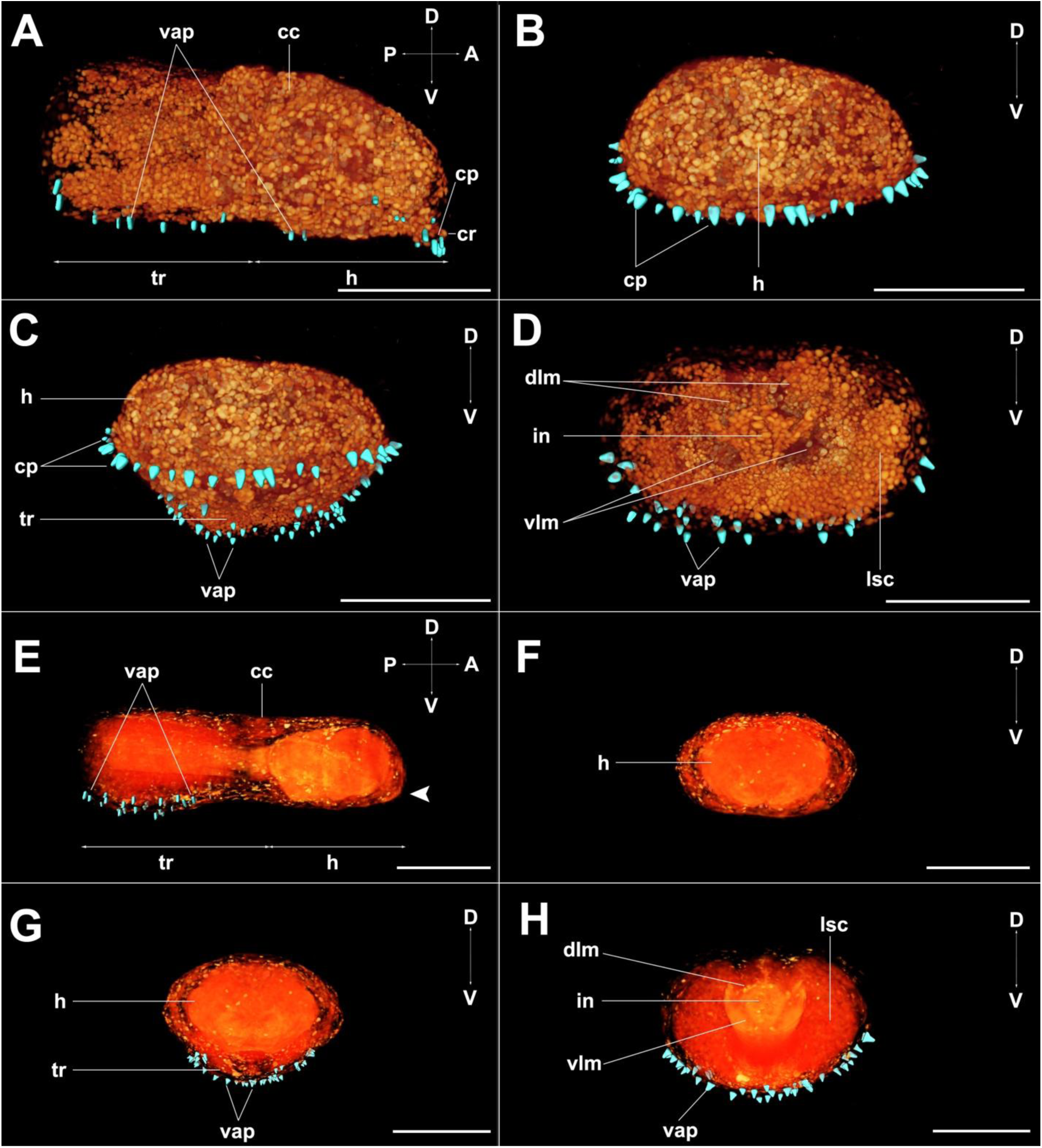
3D reconstructions of the anterior part of the body (trunk and head) of *Spadella cephaloptera* highlighting the distribution of the ventral and cephalic adhesive papillae. Volume rendering (orange) and manually segmented papillae (cyan). **A.** Lateral view in a 24 hph individual, where ventral adhesive cells (vap) are distributed in the ventral head (h) and trunk (tr) regions. Additional adhesive cells are present in the cephalic rim (cr). **B.** Front view of the anterior part of the head at 24 hph, where cephalic papillae (cp) are distributed along the cephalic rim. **C.** Tilted front view of the anterior region of the hatchling at 24 hph, where the distribution of the cells can encompass the ventral surface. **D.** Transversal section of the trunk of the hatchling at 24 hph, showing the location of the ventral adhesive papillae on the distal epidermis with respect to internal systems such as the lateral somata clusters (lsc) and the longitudinal muscles. **E.** Lateral view at 5 dph showing the absence of the cephalic rim (arrowhead) at this stage in the anterior part of the head. **F.** Front view of the anterior region of the head at 5 dph devoid of cephalic papillae. **G.** Tilted front view of the anterior region of the body, showing how the ventral adhesive cells in the trunk follow a similar distribution in the frontal plane to the one at 24 hph. **H.** Transversal section of the trunk where ventral adhesive cells form a multiple attachment system as part of the epidermis. Additional abbreviations: cc, corona ciliata; in, intestine; dlm, dorsal longitudinal muscle; vlm, ventral longitudinal muscle. Scale bars: 100 µm.

### Immunohistochemical characterization

To infer whether adhesive papillae are neuronally connected, indirect antibody staining against acetylated α-tubulin and direct staining of actin using phalloidin was performed. Strong immunoreactivity was detected in pyramidal adhesive cells (Fig. 6A-D, G-H). In the hatchling stage of *S*. *cephaloptera* (24 hph), the distribution of the microtubules related to the adhesive cells was observed in the anterior-most part of the head including the cephalic rim (Fig. 6A). These microtubules are present along the trunk and extend shortly beyond the proximities of the posterior septum (Fig. 6A). On the other hand, when reviewing the multiple tubulin processes projected from the lateral somata clusters in the VNC, no evidence was found for a direct association between individual papillae and discrete nerves (Fig. 6B). The forming cephalic ganglia do not exhibit connections with the cephalic ring papillae (Fig. 6C, D). Alternatively, a subjacent network of microtubules, related to the intraepidermal nervous plexus, is present in the distal layer of the epidermis, featuring associations with microtubules located within the cell bodies of the adhesive papillae (Suppl. Fig. 3). Additional acetylated α-tubulin^+^ and actin^+^ regions are observed in epidermal cells associated with the forming ciliary receptors (Fig. 6 A-D). The signal regarding the actin filaments within the adhesive cells was observed to be weaker across development compared to the signal from muscle-related structures (Fig. 6).

**Figure 6.**
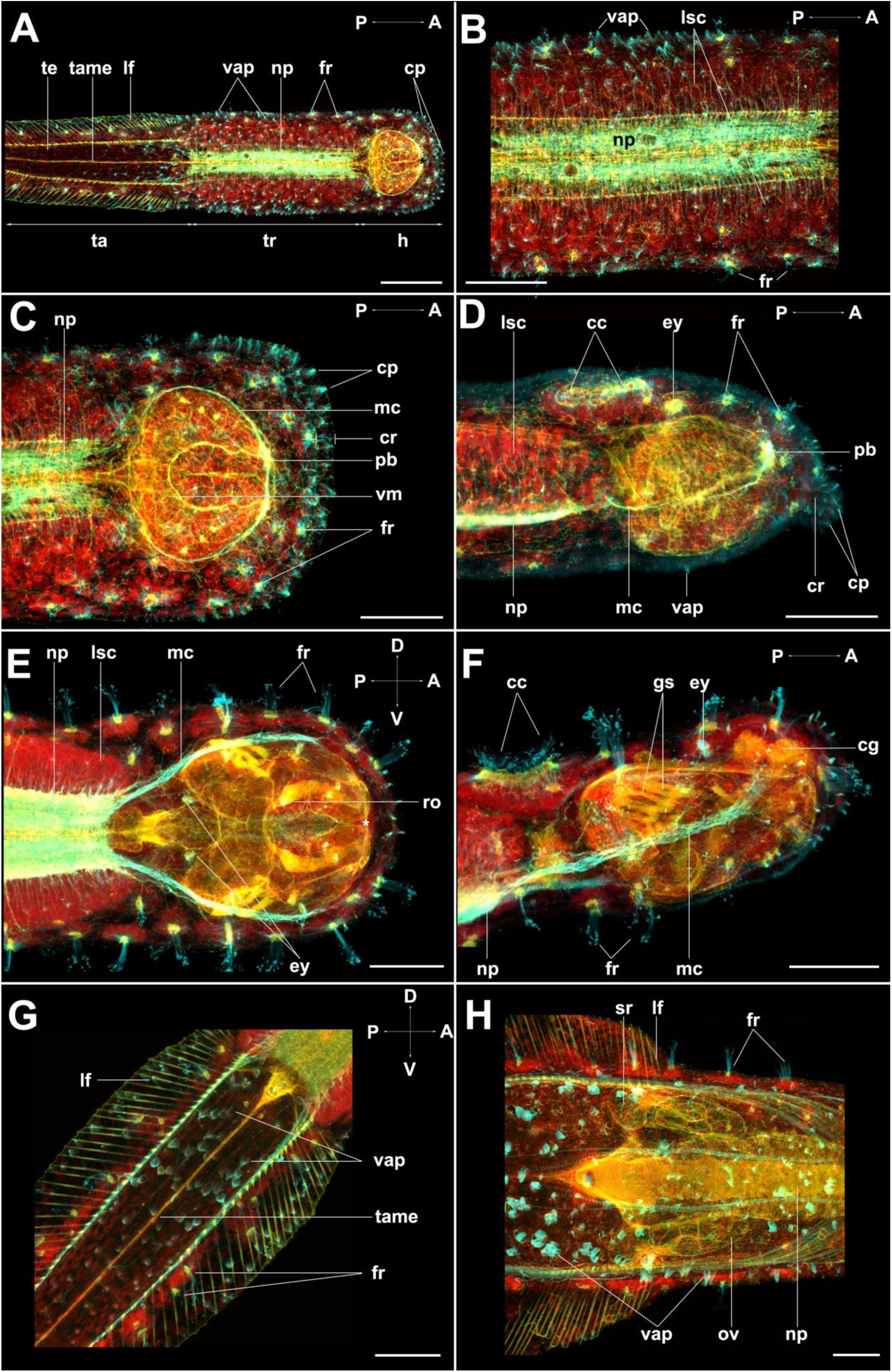
Distribution of acetylated microtubules (cyan), actin filaments (yellow), and cell nuclei (red) during the development of *Spadella cephaloptera* **A.** Overview of labelled structures at 24 hph, which include the fence receptors (fr), the ventrally distributed adhesive papillae (vap), and the cephalic papillae (cp). **B.** Maximum projection of the trunk (tr) region of 24 hph hatchling showing microtubules extending from the ventral neuropil (np), which is surrounded by the lateral somata clusters (lsc). **C.** Detail of the anterior part of the head where microtubules in the cephalic papillae distribute on the cephalic rim (cr). **D**. Side view of the hatchling, which shows the microtubule processes in the anterior end of the head. Notably, this structure is not directly connected to the primordial brain (pb). From this view, it can be seen how the fence receptors and the cephalic papillae connect to the intraepidermal plexus **E.** Maximum projection of the anterior part of the head of a 5 dph specimen, showing intense signal from acetylated microtubules associated with structures of the nervous system, such as the ventral neuropil, and the main connectives (mc). Notably, the distal epidermis is populated with multiple fence receptors, which now are in the region of the abolished cephalic rim. **F.** Side view of the head at 5 dph showing how acetylated microtubules are associated with multiple sensory organs. **G.** Detail of the tail of a 5 dph specimen showing the distribution of ventral adhesive papillae in the tail (lf)**. H.** Close up of the transition zone between the trunk and tail in the adult of *S. cephaloptera*. Additional abbreviations. cc, corona ciliata; cg, cerebral ganglion; ey, eye; gs, grasping spine; h, head; o, ovary; ro, retrocerebral organ; sr, seminal receptacle; tame, tail mesentery; vm, vestibular muscles. Scale bars: A = 100 µm; B-H = 50 µm.

At 5 dph, the specimens are distinguished by an abundance of cephalic ciliary receptors as evidenced by acetylated α-tubulin and actin processes (Fig. 6E-F). At this stage, the microtubules related to the cephalic rim identified at 24 hph were not detected. In this region, short ciliary receptors are identified (Fig. 6E). At 5dph, the adhesive papillae extend more posteriorly in the ventral region of the tail compared to the distribution observed at 24 hph (Fig. 6G). Some adhesive cells are also located on the surface of the lateral fins of 5dph individuals (Fig. 6G). Importantly, these adhesive cells are individuated and do not form clusters. Regarding the adult stage of *S. cephaloptera*, the microtubules associated with the adhesive cells are predominantly located in the anterior region of the tail, where distinct groups of clustered cells are observed (Fig. 6H). Taken together, the distribution of microtubules provides further support for ultrastructural observations of the clustering of adhesive cells happening only at later developmental stages of *S. cephaloptera* (Fig. 6H).

### Lectin affinity

To identify the location of lectin-binding glycans in whole mounts of 24 hph individuals, a panel of seven lectins was used (Figs. 7-8). Peanut agglutinin (PNA), which binds specifically to (Gal(β1– 4)GalNAc) [42], is present in the outer epidermal layer (Fig. 7A-B). In general, stronger signal was observed in the longitudinal muscles of the trunk and tail, while in the head region the vestibular musculature bordering the forming hood and grasping spines was labeled (Fig. 7A). Interestingly, PNA also strongly labels the apical region of the adhesive cells, including the papillae located in the trunk and the cephalic ring (Fig. 7B). Concanavalin A (ConA) has binding specificity for α-D-mannosyl and α-D-glucosyl groups [42]. Its binding signal was detected lining the membranes of the cells of the outermost epidermal layer across the entire surface of the hatchling (Fig. 7C). The labelling included the cell bodies of the adhesive papillae which are recognizable by their pyramidal shape of the nuclei (Fig. 7D). Soybean agglutinin (SBA) binds specifically to the terminal α and β linked N-acetyl galactosamine [42]. At 24 hph, SBA-binding signal was observed in glycans distributed throughout the epidermis, and in the posterior region lining the forming tail-coelomic cavities (Fig. 7E). In the head, SBA binding signal included the forming lateral plate, the developing eyes and the mouth opening (Fig. 7E). In addition, by reviewing the cephalic rim in the anterior part of the head, the presence of a granule-like pattern was identified, which involved both the epidermal and adhesive cells (Fig. 7F). The distribution of the binding of Ulex Europaeus Agglutinin I (UEA), which targets glycans containing α-linked fucose [42], was observed to involve the distal and proximal regions of the multilayered epidermis (Fig. 7G), extending inside the body of the chaetognath up to the proximity of the lateral somata clusters of the ventral nerve center (Fig. 7H). A closer inspection revealed that the signal inside the cell bodies of the adhesive cells is faint, suggesting a low abundance of the glycans enriched in α-linked fucose associated with UEA binding (Fig. 7G, H).

**Figure 7.**
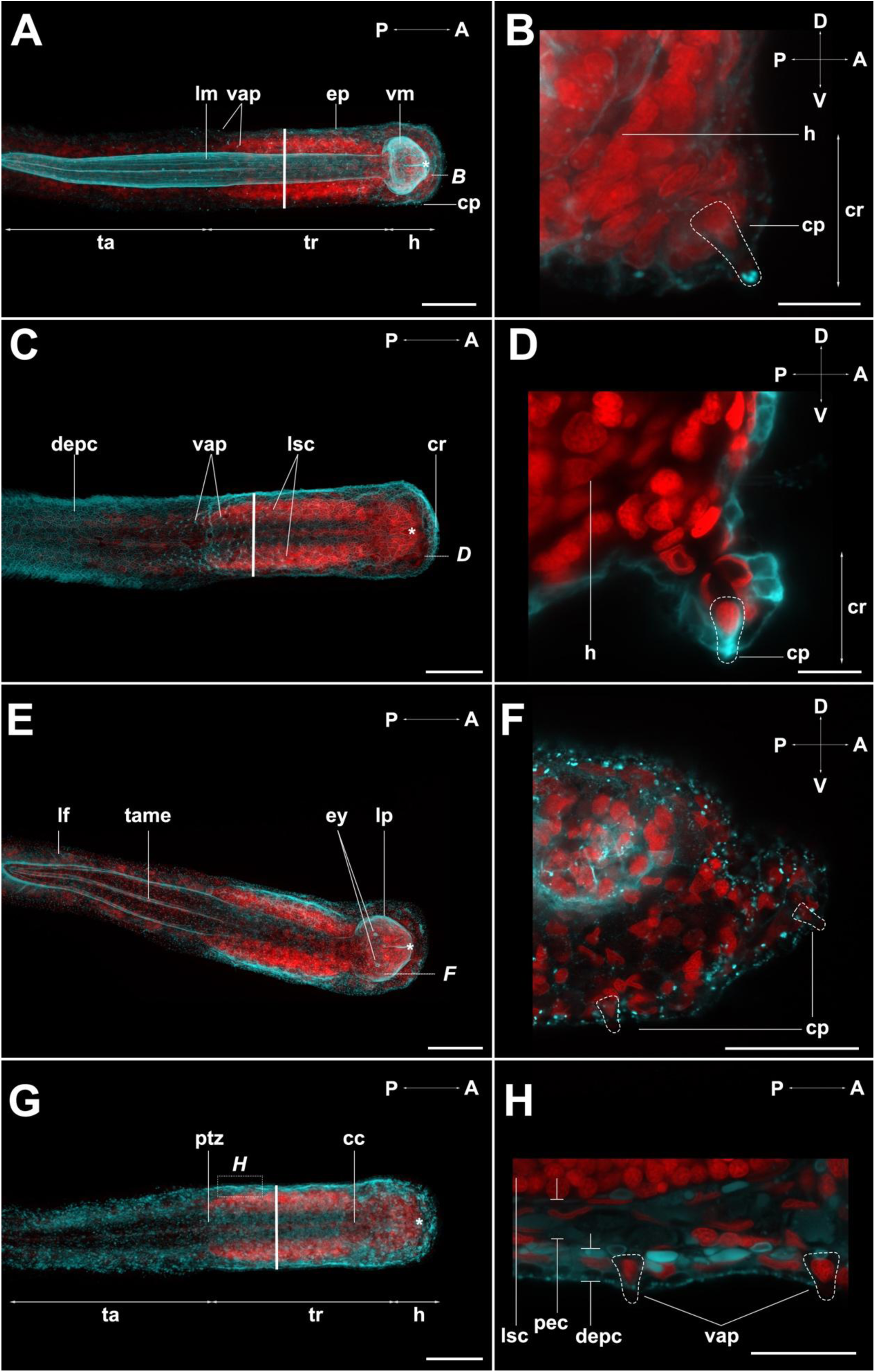
Distribution of glycans detected by lectin-binding affinity (PNA, ConA, SBA and UEA) on whole mounts of *Spadella cephaloptera* at 24 hph. The signal of the lectin histochemistry is colored with cyan, and the nuclei with red. **A.** Peanut agglutinin (PNA) related signal, labelled the longitudinal muscles (lm) and vestibular muscles (vm) and faintly the epidermis (ep) **B.** Detail of the cephalic rim (cr) in the anterior head (h) region (referenced in A), where PNA-signal strongly labels the apical region of the cephalic papillae (cp) (dotted line). **C.** Concanavalin A (ConA) bound to glycans in the cell membranes in the distal region of the epidermis (depc). **D.** Close-up of the cephalic rim (cr) (referenced in C), where glycans detected by ConA are present in the cephalic papillae (dotted line) and surrounding epidermal cells. **E**. Overview of the glycans detected by soybean agglutinin (SBA), which lined the tail coelomic cavities and the tail mesentery (tame). In the anterior region, SBA is bound to glycans in the forming lateral plate (lp) and the eyes (ey). **F.** Close-up of the anterior part of the head (referenced in E), where a granular pattern is present in both the epidermis and cephalic papillae. **G.** Maximum projection of the glycans detected by *Ulex Europaeus* agglutinin I (UEA), which are distributed on the epidermis across the body of the hatchling. **H.** Detail of the epidermis in the trunk region (noted in G), showing how UEA labeling involved both proximal (pec) and distal epidermal cells but showed faint signal in the ventral adhesive papillae (vap). Additional abbreviations: cc, corona ciliata lf, lateral fin; lsc, lateral somata cluster; ptz, posterior transition zone; ta, tail; tr, trunk. Scale bars: A, C, E, G = 100 µm; B, D = 10 µm; F = 50 µm; H = 25 µm.

**Figure 8.**
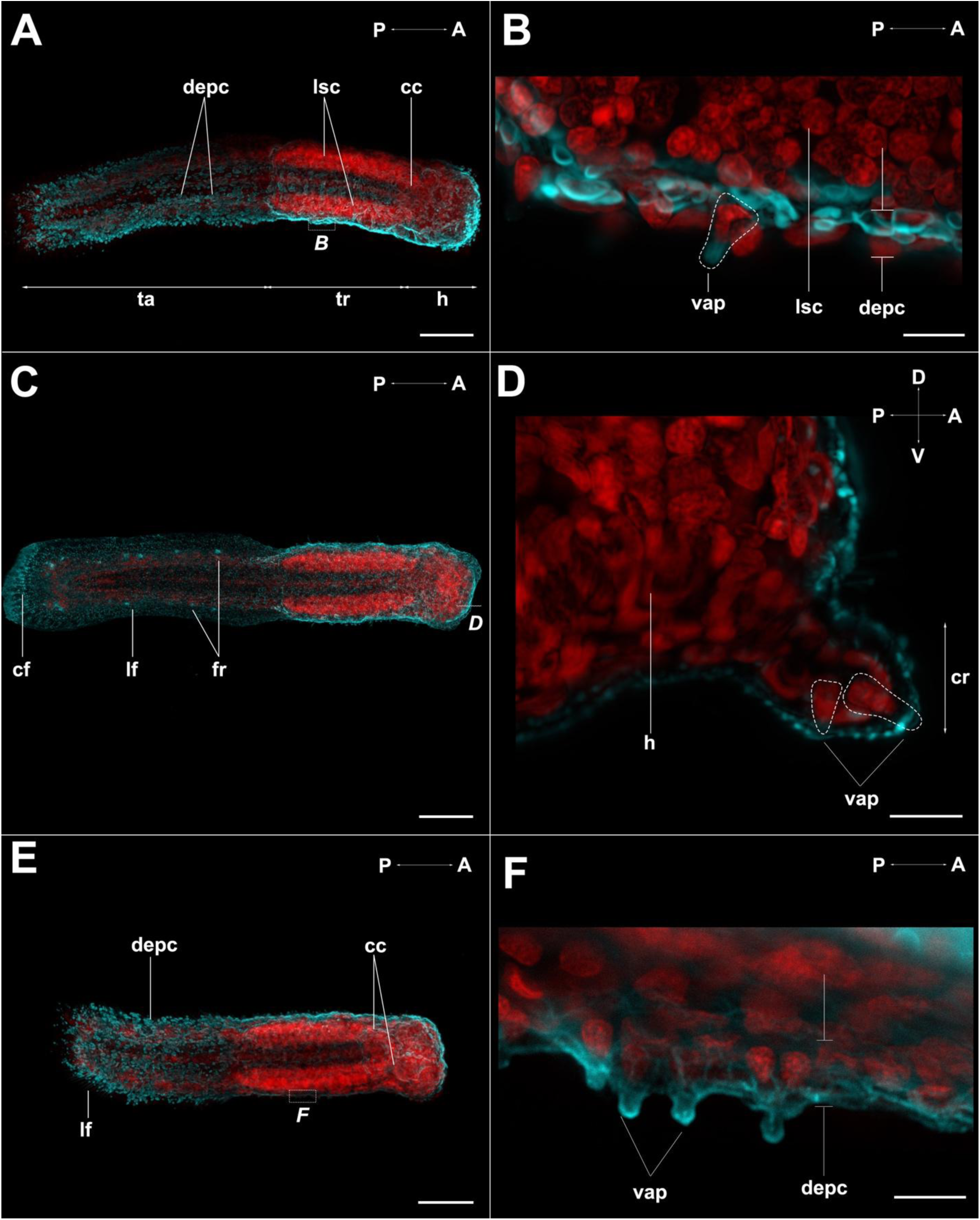
Distribution of glycans detected by lectin histochemistry (Pha-l, s-WGA and Pha-E) on whole mounts of 24 hph specimens of *S. cephaloptera*. The signal of the lectin-binding regions is colored in cyan, and the nuclei are shown in red. **A.** Overview of the glycans detected by Phaseolus Vulgaris leucoagglutinin (Pha-l), which are distributed through the epidermis. **B.** Detail of trunk (referenced in A) showing distinct labeling in the distal epidermal cells (depc) and the ventral adhesive papillae (vap). **C.** Maximum projection of the labelled glycans identified by the binding of succinated wheat germ agglutinin (s-WGA). These glycans are located in the epidermis and produce distinct labeling signal in the fence receptors (fr) distributed in the tail, and the caudal fin (cf). **D**. Close-up of the anterior part of the head (h) (noted in C), showing a granular pattern in the epidermis with no enrichment of signal in the ventral adhesive papillae (dotted lines). **E.** Location of the glycans for which Phaseolus vulgaris Erythroagglutinin (Pha-E) has binding affinity. The lectin bound preferentially to the surface of the hatchling’s epidermis. **F.** Magnified trunk region (referenced in E) showing multiple ventral adhesive papillae labelled by Pha-E, with higher signal intensity in the apical region of the cell. Additional abbreviations: cc, corona ciliata; lf, lateral fin; lsc, lateral somata cluster; ta, tail; tr, trunk. Scale bars: A, C, E = 100 µm; B, D, F = 10 µm.

*Phaseolus Vulgaris* leucoagglutinin (Pha-l) targets Galβ4GlcNAcβ6 (GlcNAcβ2Manα3) Manα3 [42]. In 24 hph specimens of *S. cephaloptera*, this moiety is present throughout the whole epidermis (Fig. 8A). Pha-l binding was observed in the two main regions of the epidermis (proximal and distal) and involved the cell body of the adhesive cells (Fig. 8B). For succinated WGA-binding glycans as chitin hydrolysate or GlcNAc with acid or salt [42], signal was observed in the ciliary receptors distributed in the tail region and with higher intensity in the caudal fin (Fig. 8C). At the same time, the location of these glycans included the entire surface of the epidermis exhibiting a granular pattern, without higher intensity inside the adhesive cells. Regarding *Phaseolus vulgaris* Erythroagglutinin (Pha-E), which preferentially binds to Galβ4GlcNAcβ2Manα6 (GlcNAcβ4) (GlcNAcβ4Manα3) Manβ4 [42], labelling was observed in the epidermis of hatchlings, strongly labelling the adhesive cells (Fig. 8E-F).

## Discussion

### Polysaccharide distribution in *Spadella cephaloptera*

Carbohydrates play an essential role in the metabolism, reproduction, structure formation, and the attachment of marine invertebrates [28, 43, 44, 45, 46, 47]. In the protandric hermaphrodite chaetognath *S. cephaloptera*, the distribution of sulfated and carboxylated polysaccharides detected by AB staining bound to anionic residues at pH 2.5 [48] was observed to be in the ciliated sperm duct, which surrounds the testis in the tail region (Figs. 1, 2, 9A-B). In adults, this duct connects the coelomic cavities, which contain the spermatogonial masses, with the seminal vesicles through a cell-ciliated funnel, as previously described by TEM observations [5]. In this view, the documented staining may correspond to carbohydrates related to glycoproteins on the surface of sperm cells, as described in the sea urchin *Paracentrotus lividus* and the tunicate *C. intestinalis*, which play a role in gamete recognition [27, 49]. However, a previous study on mature spermatozoa of *S. cephaloptera* showed that these gametes are highly concentrated in the seminal vesicles, which do not stain with AB (pH 2.5 and 1.0) (Figs. 1A, 2A) [50]. Thus, the polysaccharides are unlikely to be sperm-borne. Alternatively, the observed staining could be part of a secretion produced by the epithelial cells in the duct and be part of the seminal fluid [5]. In this sense, the epithelial cells bordering the inner side of the sperm duct, have been described to contain electron-dense secretion granules, which could be associated with the detected polysaccharides [5].

**Figure 9.**
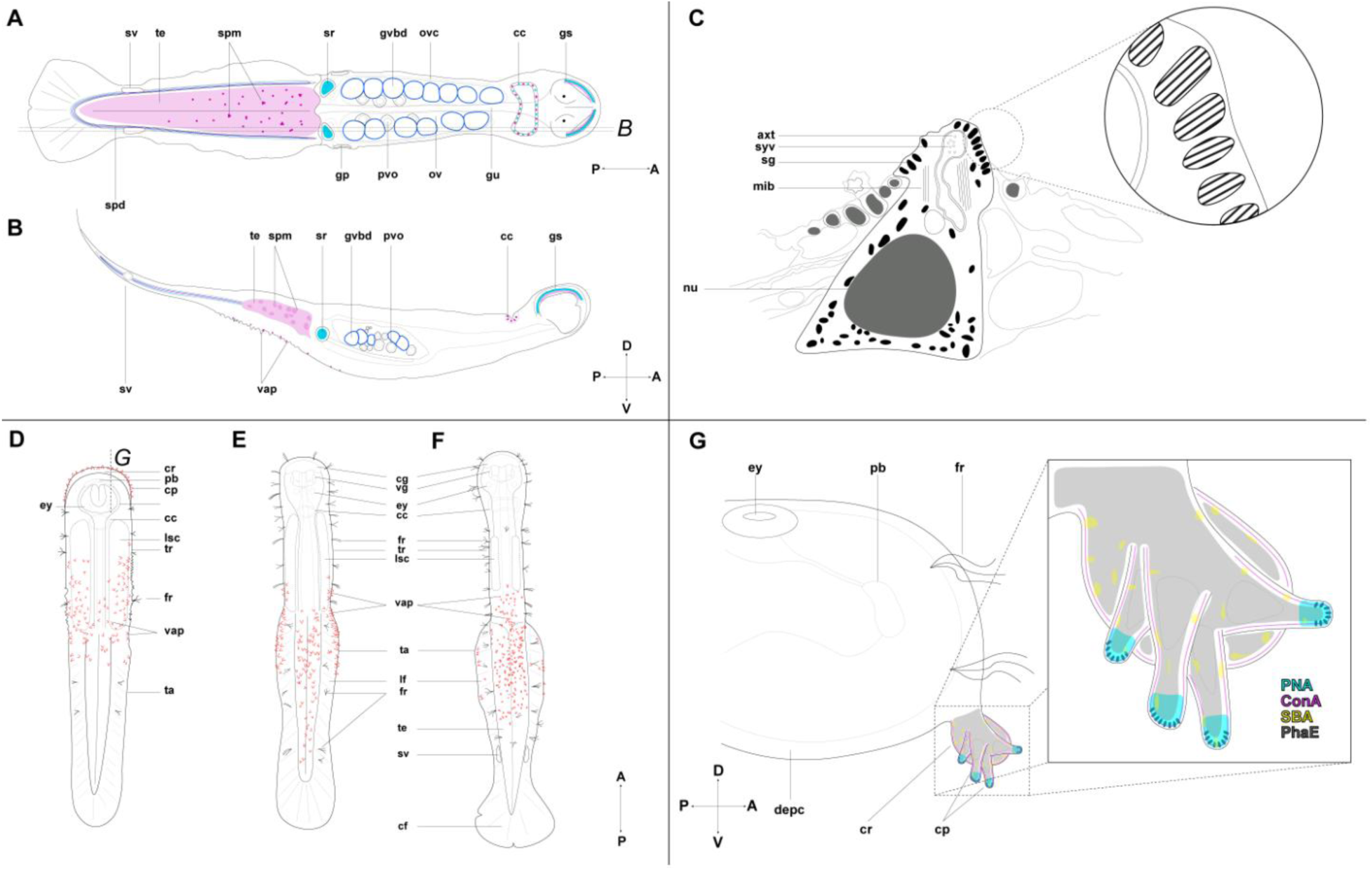
Schematic summary of the findings from the histological and ultrastructural analyses on *Spadella cephaloptera.* **A.** Distribution of glycans in the reproductive system. The location of Alcian blue (pH = 2.5)^+^ polysaccharides (dark blue) includes the sperm duct (spd) and oocytes after germinal vesicle breakdown (gvbd). Alcian blue (pH = 1.5)^+^ polysaccharides (light blue) line the sperm duct, and the seminal receptacles (sr). In the head region, the dorsal grasping spines (gs) and the corona ciliata (cc) are stained. Neutral glycans (purple) are observed within the sperm duct and in the testis (te). **B.** Sagittal section (referenced in A), showing PAS+ staining in ventral vesicle-like structures. **C.** Representation of the ultrastructure of one adhesive papilla (pyramid-like cell). These cells have secretion granules (sg), which feature fiber-like content. **D.** Ventral view of a 24 hph specimen, characterized by the presence of the cephalic rim (cr) in the anterior part of the head. This structure is enriched in cephalic papillae (cp) (red). Additional ventral adhesive papillae are in the trunk (tr). **E.** Overview of the ventral surface at 5 dph, where the cephalic rim is abolished, and adhesive cells are shifted posteriorly. **F.** Representation of the ventral view of the adult, showing the distribution of the adhesive cells. **G.** Sagittal section of the head of a specimen at 24 hph (referenced in D), where the lectins labeled moieties: PNA (cyan), Con A (magenta), SBA (yellow), PhaE (gray). Additional abbreviations: axt, axonal terminals; cf, caudal fin; cg, cerebral ganglion; depc, distal epidermis cells; ey, eye; fr, fence receptors; gp, female gonopore; lf, lateral fin; lsc, lateral somata cluster; nu, nucleus; ov, ovary; ovc, oviducal complex; pb, primordial brain; pvo, previtellogenic oocyte; spm, spermatogonial masses; sv, seminal vesicles; syv, synaptic vesicles ta, tail; vap, ventral adhesive papillae; vg, vestibular ganglion.

The presence of sulfated mucopolysaccharides has been described in the male reproductive system of diverse marine invertebrates including the sea urchin *Lytechinus variegatus*, which features such glycans in its seminal fluid forming a complex with proteins and potentially playing a role in fertilization based on the activity of sulfated glycoproteins with similar composition found in humans [51]. Alongside this, the identification of polysaccharides in the seminal fluid of marine invertebrates has been observed to be key for the survival of the male gametes. For instance, the sperm cells of the crab *Scylla serrata*, exhibit higher activity of enzymes related to carbohydrate metabolism under anaerobic conditions and feature a decrease in polysaccharide concentrations when they are contained in the female organ, which is usually devoid of this substrate [52]. Considering the above, it is possible that the polysaccharides in the seminal fluid of *S. cephaloptera* may be used to favor the survival of the sperm cells when they are released and subsequently stored in the oviducal complex before fertilization [26].

A comparable distribution of carbohydrates to that reported here for *S. cephaloptera* has also been demonstrated in distant taxa, such as in the arthropod *Goniopsis cruenata*, for which secreted polysaccharides with an epithelial origin were observed inside the vasa deferens using AB [53]. It was proposed that these secreted polysaccharides could be involved in the development of the sperm cells [53]. Together, our data for *S. cephaloptera,* revealed the presence of sulfated polysaccharides in the sperm duct (Figs. 2A-B, 9A-B), hinting at a hypothetical conserved mechanism among marine invertebrates that utilizes the seminal fluid enriched in fucoidal polysaccharides to aid sperm cells in reaching their destination inside the ovaries. Moreover, the distribution of carboxylated and sulfated polysaccharides, showed that these two types of mucopolysaccharides are distributed in the sperm duct of *S. cephaloptera* (Figs. 1C, 2A-B, 9A-B). Either both types of polysaccharides are present in the duct as observed in the gastropod *Haminoea navicular* [54], or a single polysaccharide with mixed charge (e.g., chondroitin sulfate) is detected at both pH levels [55]. Such a pattern was observed in the dendrobranchiate crustacean *Penaeus indicus,* which features hyaluronic acid (carboxylated) and chondroitin sulfate (carboxylated and sulphated) in the spermatophore [56]. Further chemical characterizations of the seminal fluid would be necessary to disentangle the identity of the individual mucopolysaccharides in the sperm duct of *S. cephaloptera*.

The distribution of neutral polysaccharides visualized by the PAS^+^ reaction in *S. cephaloptera* was localized in the sperm ducts in the tail region (Figs. 3A, 9A-B), which are lined by ciliated and secretory epithelial cells [5]. In this sense, the neutral glycan observed in the vas deferens *of S. cephaloptera* could be also part of an epithelial secretion as suggested for the alcianophilic content of this cavity. This distribution of neutral polysaccharides in the vas deferens has also been reported in the crustaceans *Eriphia verrucosa* and *Callinectes danae*, where the secretory epithelium of this structure is surrounded by a PAS-positive coelomic fluid [57, 58]. In *S. cephaloptera*, additional staining is visible in the coelomic cavities of the tail (Fig. 3A), which could correspond to glycogen, a neutral polysaccharide detectable by the PAS reaction in the forming sperm cells within the spermatogonial masses. A similar distribution of neutral PAS^+^ polysaccharides in spermatogonial cells and forming spermatozoons has been reported in the shrimp *P. indicus* [56]. Likewise, it has been demonstrated that stored glycogen concentrated in the flagellum of the sperm is a widespread feature present in different taxa, including annelids, mollusks, echinoderms, and chordates [59, 60, 61] The presence of this polysaccharide has been proposed to serve as metabolic fuel for the energetic requirements of the spermatozoa during the fertilization process, and it can represent a significant proportion of the composition of the whole gamete as is the case of the tunicate *C. intestinalis*, where it constitutes ∼3% of the sperm dry mass [59]. Accordingly, the detection of this glycan in the male reproductive structures hints at the idea that *S. cephaloptera,* among others may use this glycogen-dependent, evolutionarily conserved mechanism to provide metabolic resources to its sperm cells.

Additional roles of mucopolysaccharides in marine invertebrates include mediating recognition mechanisms between oocytes and sperm cells, which are essential to ensure species-specific fertilization [62, 63]. In the case of *S. cephaloptera*, there is evidence of a jelly coat being present from the vitellogenic phase of the largest oocytes that will be involved in fertilization [5, 26].

Nevertheless, chaetognaths are characterized by a unique fertilization mechanism in which direct interactions between the sperm cells and the jelly coat do not take place [26, 64]. The sperm cell reaches the oocyte through a fertilization canal built by accessory fertilization cells (AFC) [26, 64]. In the present study, it is shown that the jelly coat contains carboxylated polysaccharides detected by AB (pH = 2.5) (Figs. 1E-F, 9A-B), however no evidence of sulfated polysaccharides as revealed by AB (pH = 1.0) staining exists (Figs. 2C, 9A-B). Considering these features, the polysaccharides present in the coat could act as ligands for receptors in the ACF, potentially ensuring that only mature oocytes are exposed to the sperm in the oviducal complex. These observations contrast with what is observed in the sea urchin, where there is evidence that sperm activation is triggered by direct molecular interactions with fucosylated sulfated polysaccharides in the surface of the egg coat [63, 65]. On the other hand, the carboxylated nature of the polysaccharide covering the mature oocytes in *S. cephaloptera* follows the chemical categorization of non-sulfated chondroitin, which is also present in the oocytes of *Caenorhabditis elegans,* being a key agent in fertilization and development [66, 67]. However, there are few reports of non-sulfated mucopolysaccharides in marine invertebrates, except for the one involved in the tubulation of the nematocyst in *Nematostella vectensis* [68]. Therefore, despite the preliminary characterization of these glycans presented here, further chemical characterization is necessary to determine the identity of the molecules present in the female reproductive system of *S. cephaloptera*.

The presence of PAS^+^ vesicles in the ventral clusters of papillae in the posterior region of the epidermis of *S. cephaloptera* (Figs. 3A-B, 9B) suggests that these epidermal cells, which are thought to mediate the attachment process [5, 13], potentially secrete neutral hexose mucosubstances (Histological and histochemical methods). This type of mucosubstance has been described in multiple temporary attachment systems, with extensive reports on different cephalopods. For instance, in the cephalopod *Euprymma scolopes*, the adhesive gland cells (goblets) located in the dorsal epithelium contain neutral sugars [69]. In the case of *Nautilus pompilius*, neutral polysaccharides form PAS^+^ granules within the glandular cells of the tentacle [70]. Additionally, two species of *Idiosepius spp.* exhibit neutral carbohydrates in the columnar and granular cells located in the distal epidermis of the adhesive organ [71]. Representatives from other taxa featuring this trait include the hydrozoan *Hydra vulgaris*, in which glycans detected by PAS were found in the basal disc, as well as secretory granules [72]. In this view, the concentration of PAS^+^ glycans in the papillae of *S. cephaloptera* suggests that secretions of these cells with neutral mucopolysaccharide vesicles could be involved in the attachment process, as it appears to be the case for many other marine invertebrates.

On the other hand, the presence of neutral polysaccharides has also been documented in marine invertebrates that exhibit permanent attachment. In the barnacle *Lepas anatifera*, the cytoplasm of the adhesive gland cells and the principal canal that transports the glue to the substratum both exhibit neutral mucopolysaccharides [73]. Interestingly, there is evidence for *Latia neritoides*, a fresh-water living mollusk showing PAS+ molecules in both its adhesive cells and secretion [74]. In this context, the widespread presence of neutral glycans supports the notion of conservation at the molecular level of marine attachment systems [16]. Along with this, it has been previously proposed that the presence of glycans provides support through electrostatic interactions of the polar groups present in these molecules and the surfaces of contact, which could explain their prevalence across distant taxa [2, 75].

### Ultrastructure of the adhesive cells

The ultrastructure of the adhesive cells of *S. cephaloptera* revealed the conformational changes the attachment system undergoes during development. From hatching up to 5 dph, individuals of *S. cephaloptera* exhibit discrete, individuated adhesive cells embedded in the distal layer of the epidermis, which also hosts numerous electron-lucent vacuoles and secretory granules (Figs. 4A, 9C) [5]. During ontogeny, the cells of the adhesive epithelium are swollen in the tip and have a pyramidal-like shape (Figs. 4A, C, E, 9C), a feature observed in other members of Spiralia, such as the cephalopod *E. scolopes* [76]. In the early stages of development of the hatched *S. cephaloptera* specimens, the epidermal cells surrounding the adhesive papillae are active in terms of merocrine exocytosis (Fig. 4A) [5] and form an epidermal layer with a thickness of ∼5 μm. In contrast, in late juveniles and adults of *S. cephaloptera* this tegument layer was observed to grow up to ∼7 μm and maintains this secretory activity. Notably, during these stages of late development, the adhesive cells form clusters and are in physical contact with each other inside these cell groups (Fig. 4E). The presence of microtubules was recognized in the adhesive cells (Fig. 4B, F), which could provide resistance against displacement, such filament support has been also observed in the anchor cells of *Macrostomum lignano*, playing an essential role in the attachment [18]. Another feature present across different attachment systems is a close relationship to the nervous system. For instance, in the case of *A. rubens*, independent neurosecretory cells display a close association with the nerve plexus [77]. A neuronal regulation of secretory granules is also suggested in the cases of *Idiosepius biserialis* and *Idiosepius pygmaeus,* where fusiform cells respond to stimuli and induce the secretion of the adhesive substance [71]. In *S. cephaloptera*, the presence of synaptic vesicles in the apical region of the adhesive cells (Figs. 4B, D, F) [5], together with previous TEM evidence of basiepidermal neuronal plexus processes [5], provides support to the idea that the nervous system of the chaetognath plays a role in the attachment process.

During ontogeny, electron-dense secretion granules of about 240 nm in diameter were observed in the adhesive papillae of *S. cephaloptera*. Furthermore, these secretory products were abundant in the apical part of the cell (Figs. 4A, C, E, 9C). A similar distribution pattern has been observed in the adhesive appendages of the close relative chaetognath *Paraspadella gotoi,* which suggests that these systems may favor polarized storage as it would allow rapid release upon specific stimuli [78].In adult specimens of *S. cephaloptera* we observed that these electron dense granules can presumably fuse with the cell membrane and release their content (Supp. Fig. 3). Consequently, the ventral papillae across different developmental stages of *S. cephaloptera* supposably behave much like exocrine glands that undergo regulated exocytosis [5]. The presence of secretion granules in non-permanent attachment systems is a widespread trait observed in several representatives from different taxa such as mollusks and flatworms [17, 79]. Specifically, TEM analysis of the adhesive cells of *S. cephaloptera* (Figs. 4D, F, 9C) revealed, for the first time, the fibrous nature of the content inside the secretion granules produced by those cells. Similarly, previous research on the ultrastructure of the adhesive appendages of the chaetognath *Paraspadella gotoi* featured secretory granules with two distinct electron dense regions, which exhibited parallel strands in their cross-sections [78]. In addition, this type of organization within the secretion granules has also been described in members of different taxonomic groups. For instance, in the sea urchins *Arbacia lixula* and *Sphaerechinus granularis,* the granules exhibit parallel plates [79].

Nevertheless, although the similarities found with chaetognaths, it has also been demonstrated that in echinoderms, the secretions granules can display species-specific features regarding their shape and organization of their content [79]. Additionally, the formation of parallel plates has been previously hypothesized in the adhesive systems of marine invertebrates to be linked to the presence of functional amyloid proteins [80, 81]. In this sense, it is known that amyloid-like proteins can form extensive fibers within the cell bodies due to the stability provided by the β-sheet arrangement, which might play a role in the attachment process and stabilization of the ECM [82]. A recent study highlighted the high prevalence of marine adhesive proteins featuring epidermal growth factor (EGF)- like domains, which are beta-sheet forming regions that may confer stability to those proteins [3]. In the case of *S. cephaloptera*, transcriptomic cell-type analyses may help clarify the molecular identity of the fibrous content in the secretion granules of the adhesive cells.

Another widespread feature of some marine temporary attachment systems is the presence of detachment cells coupled to the adhesive cells, forming a dual-gland system. Such systems are present in a variety of taxa, including flatworms and sea stars [83, 84]. In *S. cephaloptera,* as observed in the TEM images here presented (Fig. 4), the adhesive cells are not surrounded by specialized or distinct cells, but instead by regular epidermal cells (Figs. 4A, C, E, 9C). There is evidence that some marine invertebrates do not rely on additional secretions for detachment, but rather on muscle contractions, proposed for the detachment of *Euprymna scolopes* [16, 76]. In the context of chaetognaths, *P. gotoi* has dedicated muscles (supercontractors) linked to the adhesive appendages that allow a range of movements that include swings and jumps when contracted [78]. Remarkably, these muscles have been reported across members of the genera *Paraspadella spp.* and *Spadella spp.* [5, 78]. Additionally, the behavior of *S. cephaloptera*, which does not involve constant swimming but rather quick flicks [13], would require a fast-reaction detachment mechanism. Therefore, it is feasible that in *S. cephaloptera*, supercontractor muscles could be involved in the detachment process, as they do in other benthic species [78].

### Spatial distribution of adhesive cells across development

The location of the adhesive cells across the different developmental stages investigated exhibited changes that matched the behavior physiology of the chaetognath (Figs. 5, 9D-F). In this regard, at 24 hph individuals of *S. cephaloptera* do not have a developed mouth opening, a digestive track, or a functional anus that would allow them to feed or digest food [11]. For these reasons, they do not lift their heads to catch prey and keep the anterior part of the body attached to the surface they are in contact with [5, 13]. Observations from a previous study indicated that after hatching, specimens of *S. cephaloptera* remain in the vicinity of the region where the eggs were laid and form groups [13]. This remark can be explained by the presence of adhesive papillae in the cephalic rim, which keeps the hatchling firmly attached (Figs. 5 A-C, 9D). The distribution of elongated adhesive cells in the anterior part of the head during early stages appears to be a conserved trait among other members of Spadellidae. For instance, in the case of *Spadella schizoptera,* evidence of cephalic papillae is observed up to the fourth day of development after hatching [85]. Similarly, we found that individuals of *S. cephaloptera* at 5 dph do not have cephalic adhesive papillae (Figs. 5 E-G, 9E). In this context, previous research on hatchlings of *S. cephaloptera* showed that between 5dph and 7dph the alimentary canal and anus form, which suggests that at this moment during ontogeny the animal starts to feed, as the yolk reserves deplete [5, 13]. The present study provides support for this view, as the absence of adhesive cells in the anterior region (Figs. 5E-G, 9E), enables the animal to promptly lift its head and capture prey. Consequently, the maturation of structures such as the esophageal ganglion, which is thought to target esophageal muscles through the esophageal nerve, and vestibular ganglia, presumably controlling the grasping spines, likewise progress towards a higher degree of complexity after between 72 and 96 hph [5, 11]. A key point is that in the case of *S. schizoptera,* there is no evidence of ventral papillae being present in the trunk during development, but rather the formation of a dedicated adhesive organ in the tail [85].

On the other hand, our observations in *S. cephaloptera* show several ventral papillae distributed along the trunk, establishing a multiple attachment system as described for other metazoans (Figs. 5A, E, 9D-F) [16]. At 5 dph, additional adhesive cells located ventrally in the most anterior part of the trunk and the head at 24 hph (Figs. 5 A, C, 9D), are no longer present (Figs. 5 E-G, 9E), which provides support to the idea that over development individuals of *S. cephaloptera* need dynamic lifting of the anterior part of their bodies. In this view, the distribution of the cells *S. cephaloptera* during ontogeny ensures a stable habitat shortly after hatching but shifts once the animal relies on hunting for feeding, which coincides with abolition of the transient cephalic adhesive papillae.

### Actin and microtubule distribution in the attachment system

The distribution of acetylated microtubules has been previously investigated in *S. cephaloptera* mainly to study the features of its nervous system [11]. While Rieger et al. (2011) demonstrated the immunoreactivity of the ventral adhesive cells in the overview of different developmental stages, the present study searched for direct connections between the different neuronal ganglia and the adhesive cells (Fig. 6). In this aspect, no distinct tubulin-positive nerve processes were found projecting directly from the ventral nerve center (Fig. 6B) or the cephalic ganglia (Fig. 6C) to individual adhesive cells. Nevertheless, during ontogeny it was observed that the multiple acetylated microtubule bundles inside the cell bodies of the adhesive papilla projected into the epidermis, known to host an intra- and basiepidermal neuronal plexus, which is part of the peripheral nervous system and is considered an autapomorphy of chaetognaths [5, 11]. Moreover, as shown in Fig. 6C, the different mechanosensory receptors (ciliary fence and tuft receptors) embedded in the epidermis of *S. cephaloptera* also display enriched abundance of acetylated microtubule fibers that protrude outside while anchoring to the tegument. In the epidermal region, a previous study suggested that these microtubules are connected to the intra- and basiepidermal plexus, which communicates input sensed by the different ciliary receptors located through the body of *S. cephaloptera* to the VNC [5]. In addition, previous TEM observations from adults of *S. cephaloptera* have pointed out that there are condensed neurons and neuronal somata below the distal epidermis layer, which also hosts the adhesive cells in *S. cephaloptera* [5]. Given the above and considering the evidence of acetylated microtubules that connect the ventral adhesive papilla to the intraepidermal neuronal plexus [11] (present study), the adhesive cells might or provide input to the VNC and could participate in a neuro-secretion mechanism that releases on demand the secretion granules containing agents potentially involved in the attachment process. This hypothesis is further supported by the evidence of synaptic vesicles in the cell bodies of the adhesive cells observed through the ontogeny of *S. cephaloptera* (Fig. 4B, D, F) [5]. The role of microtubules in adhesion has also been documented in *C. intestinalis* and *Ascidia malaca,* where the signal is localized in the adhesive cells, hinting at a sensory function of these cells [86]. The idea of the innervation of the attachment system of *S. cephaloptera* is also supported by the fact that the animals are unable to attach during relaxation using MgCl_2_ before fixation (data not shown). MgCl_2_ is known to compete with ions in the synapses of nerves and muscles [87]. An additional role of acetylated microtubules would be to provide mechanical support for the adhesion cells, as it has been described that acetylation of microtubules increases resistance and features higher stability [88]. Considering these findings, the distinctive distribution of acetylated α-tubulin in the ventral papillae suggests the capacity of these cells to communicate with the nervous system and potentially provide additional structural support for attachment.

Here, we show for the first time, that the distribution of F-actin filaments in *S. cephaloptera* is enriched at the base of the ciliary receptors, probably providing support for the structures that project outside the body (Fig. 6C, E, F). Conversely, in the adhesive papillae, the presence of F-actin bundles was lower but still detectable in the cytoplasm of these cells (Fig. 6C, G). The relevance of actin filaments in the attachment systems of marine invertebrates has been demonstrated in *Macrostomum lignano*, where anchor cells display microvilli with actin-dense cores under normal conditions and a reduced density of bundles when mutations in anchor cell proteins are induced [84]. Nevertheless, the low level of intensity of the actin filaments in the ventral papillae of S. *cephaloptera* hints towards the idea that the attachment relies on additional mechanisms for structural support.

In comparison to other members of Spadellidae, *S. cephaloptera* exhibits an attachment system that displays core conserved traits while also featuring noticeable differences from the adhesive systems documented in other members of this family. For instance, during early development, *S. schizoptera* also exhibits a cephalic rim of elongated adhesive cells in the anterior part of the head; however, the presence of this structure is reported up to 4 dph, suggesting that the cephalic attachment is also abolished during ontogeny [11]. On the other hand, at 5 dph in *S. cephaloptera,* individuated papillae were observed (Fig. 6G), which in adulthood form clusters of cells distributed in the tail region (Fig. 6H). Interestingly, adult specimens of *S. schizoptera* exhibit different, more complex structures for attachment in the posterior part of the body, relying on a finger-like adhesive organ with multiple digitations covered with densely aggregated and less elongated papillae [85]. Another chaetognath, *Spadella japonica,* exhibits what seems to be an intermediate phenotype between *S. cephaloptera* and *S. schizoptera*, featuring papillae in the antero-ventral tail region, and next to them two small processes covered with adhesive papillae [89, 90]. This attachment organ resembles the shape of one of the digitations of *S. schizoptera,* covering the adhesive papillae that approximate in shape those observed in *S. cephaloptera*. Notably, *S. japonica* displays a network of adhesive papillae in the adult stage as observed using SEM, which parallels the observed distribution in the early stages of *S. cephaloptera* [89]. However, there are no TEM or immunohistochemistry data that indicate whether all the observed papillae in this species are individuated or could form clusters of cells, as in the adult of *S. cephaloptera*.

### Distribution of lectin-binding glycans in the hatchling stage

Glycans have been demonstrated to be part of different temporary attachment systems in marine invertebrates through results obtained from histological stainings (e.g., PAS staining, AB) [69] and immunohistochemistry based on the affinity of lectins for specific moieties [18, 19, 79, 91, 92]. Likewise, it has been described that specific glycans are linked with proteins located in the attachment organ. For example, in the sea urchin *P. lividus*, lectins known to target N-acetylglucosamine exhibited a specific binding affinity for proteins extracted from the tube feet when these were immunoblotted [93]. This approach demonstrated the presence of N-glycosylation in the adhesive proteins, a post-translational modification produced during protein folding that reduces dynamics and potentially adds stabilization [94]. In the present study, we demonstrate the binding affinity of PNA (Peanut agglutinin) to the most apical part of the adhesive cells of *S. cephaloptera*, producing an intense signal (Figs. 7B, 9G). In this region of the cell body, as observed by TEM, there is higher concentration of the electron dense granules (Figs. 4A, C, E, 9G). PNA is known to bind the covalently attached serine/threonine O-linked glycans [95]. In this sense, O-glycosylation of human proteins occurs in folded proteins [96], and has been demonstrated to increase their global stability while reducing flexibility [97]. Interestingly, this type of post-translational modification has also been linked to favor protein secretion [97]. From these observations, it is suggested that either the secretion granules contain a heavily O-glycosylated protein, due to the strength of the lectin signal, or some galactose-containing carbohydrate [98], which may be sufficiently stable when secreted to potentially mediate interactions for the attachment of *S. cephaloptera*.

The affinity of PNA for adhesive cells of marine invertebrates has been described in diverse taxa. For instance, in lophotrochozoans, such as the platyhelminth *Macrostomum lignano*, PNA produces specific binding to the adhesive cells and the rim of the electron-dense core vesicles [18]. Notably, there are reports of freshwater organisms, such as *Schmidtea mediterranea*, for which PNA has also been described to bind specifically to gland cells [99]. In the sea star *Asterina gibbosa,* PNA labelled the gland cells, where the binding pattern resembled elliptic dots [17]. Notably, results from the echinoderm *Asterias rubens* showed the presence of lectin binding regions of proteins in the footprint (including PNA), implying that O-glycosylation plays a role in the attachment process [2]. On the other hand, for deuterostomes, the labelling by PNA included the collocytes of the tunicate *C. intestinalis* [19]. Overall, the binding of PNA underlies a widely distributed feature of different marine temporary attachment systems, which, based on the results of lectin blots using footprint proteins, is related to O-glycosylation [2, 100]. In this view and considering the labeling distribution of PNA in *S. cephaloptera*, we hypothesize that this type of O-glycosylation may be part of the convergently evolved features that several marine adhesive proteins exhibit [16].

The lectin Concanavalin A (α-Man/α-Glc-binding lectin) strongly labelled the membranes of the epidermis and the ventral papillae of *S. cephaloptera* (Figs. 7C-D, 9G). A similar labelling pattern has been previously reported in the epidermis of *M. lignano* [18]. Likewise, it has also been described as part of the footprint of *A. rubens*, *A. gibbosa,* and weakly labeling the adhesive secretion of *P. lividus*, suggesting a role of these glycans in the attachment process of echinoderms [2, 17, 93]. Furthermore, the presence of ConA-binding glycans is consistent with the presence of N-glycosylated proteins occurring inside the adhesive cells [101]. Moreover, it has been proposed that functional amyloids are related to the underwater properties of the marine adhesives [82]. Recent research has shown that N-glycosylation in other eukaryotes limits protein aggregation prone to misfolding [102]. In this view, it is plausible that, besides providing electrostatic degrees of freedom for the glue, these post-translational modifications may help prevent uncontrolled aggregation of the protein content before its release. In *S. cephaloptera*, there is no confirmation of amyloid forming structures. However, the content of the secretion granules exhibits a fibrillar conformation (Fig. 4D, F); thus, thioflavin staining may clarify whether the fibrils possess amyloid properties.

In the case of the lectin SBA, the distribution of labeling involved both the epidermis and the adhesive cells of *S. cephaloptera*, resembling a granular pattern with reduced specificity and abundance in the adhesive papillae compared to PNA and ConA, respectively (Figs. 7E, F, 9G). Interestingly, in *A. gibbosa*, a similar pattern was observed in the disc epidermis, where elliptical dots indicated the presence of N-acetylgalactosamine [17]. Additional evidence of SBA-binding glycans has been confirmed in the adhesive organ of *P. lividus* using lectin histochemistry [93]. This lectin has also been used for the confirmation of glycoproteins containing the O-linked N-acetylgalactosamine in the adhesive disc of *P. lividus* through lectin pull downs [100]. In *M. lignano*, there was no indication of the binding of SBA to the adhesive organ cells [18]. On this basis, it can be noted that the N-acetylgalactosamine monosaccharide has less specificity for the previously investigated adhesive organs of marine invertebrates when compared to the forming disaccharide detected by PNA. This observation can be related to the fact that PNA-binding glycans have an ubiquitous presence on mucins, which have been reported to be part of the adhesive systems of marine invertebrates [14]. Moreover, high levels of O-glycosylation have been linked to polymeric gel-forming mucins with the capacity to form hydrogels, which would allow for maintaining contact with the substrate while preserving the viscoelastic properties of the marine glues [103].

The labelling pattern of UEA, which binds to αFuc-linked on galactose residues, is of low intensity in the cell bodies of the ventral papillae of *S. cephaloptera* (Fig. 7H). The presence of UEA-binding glycans in other temporary attachment systems from marine invertebrates is scarce, being detected strongly only in lectin blots from the proteinaceous fraction of the adhesive material of *A. rubens* [2]. Therefore, this type of glycan appears to play a more species-specific role. Regarding the moieties detected by Pha-L-binding, such as Galβ4GlcNAcβ6 (GlcNAcβ2Manα3) Manα3, they were observed in the epidermis and the ventral papillae of *S. cephaloptera* (Fig. 8A, B). Comparable results have been reported in the adhesive organ and footprint of *M. ileanae,* and with moderate intensity in the adhesive disc of *P. lividus* [32, 93] but absent in *C. intestinalis* suggesting that they are also part of less conserved mechanisms [19].

In the case of succinated WGA, known to bind GlcNAc, no papillae-specific signal is evident in *S. cephaloptera* but instead, a granular pattern can be observed in the distal epidermis (Fig. 8C, D).This lectin has been found to bind to the cells of multiple attachment systems, including representatives from tunicates and echinoderms, but not described in the adhesive organ of plathelminths [18, 93]. On the other hand, the lectin PHA-E, which has affinity for multiple sugar moieties (Galβ4GlcNAcβ2Manα6 (GlcNAcβ4) (GlcNAcβ4Manα3) Manβ4), and in *S. cephaloptera*, it strongly labelled the distal layer of the epidermis (Fig. 8E). The signal was also observed in the cell bodies of the ventral papillae, with the brightest signal at the cell apex (Figs. 8F, 9E). A similar distribution has been reported across tunicates including the larvae of *C. intestinalis*, where PHA-E bound preferentially to glycans in the tip of the papillary organ, and the adhesive plaque [19]. On this basis, even though the GlcNAc residue is part of the target of multiple lectins, when it is investigated in isolation, there is no evidence that it occurs with higher prevalence in the ventral papillae of *S. cephaloptera*. On the other hand, additional moieties present in the glycans such those in Galβ3GalNAc (targeted by PNA), and GlcNAcβ4Manα3 (targeted by PHA-E), are enriched in the adhesive cells, suggesting that more complex glycosylation processes may be involved in the conformation of these cells, and consequently in the attachment process of chaetognaths.

## Conclusion

The present study applied histochemistry, lectin labeling, transmission electron microscopy, and cytoskeletal mapping and reveals that the chaetognath *Spadella cephaloptera* uses glycan-rich epithelial secretions and not sperm-borne glycans in its reproductive tract. Furthermore, neutral glycan-containing vesicles were observed in the adhesive cells. These cells also feature apical electron dense secretion granules that exhibit fibrillar content that likely mediates the attachment. Moreover, the adhesive papillae are restricted to the anterior cephalic and trunk region of hatchlings. During subsequent development they degenerate in the cephalic region, but the trunk and tail regions are enriched in these structures. Additionally, the adhesive papillae exhibit traits such as synaptic vesicles and microtubule associations with the intraepidermal plexus, indicating a neurosecretory component. Considering the above, the attachment system of *S. cephaloptera* combines species-specific glycan signatures (O- and N-linked moieties) with hallmarks of convergently evolutionary features of temporary adhesive systems.

## Declarations

### Ethics approval and consent to participate

The research performed on chaetognaths does not require either ethics approval or consent to participate.

### Consent for publication

Not applicable

### Author contributions

CB and TW conceived the idea and planned the project. TW secured funding for the project. CB collected the specimens. JB and ST standardized the fixation for TEM. JB performed histological staining experiments. CB, ST and JB produced the resin/paraffin blocks and performed the sectioning for the data acquisition. ST and JB prepared the samples for TEM. JB supervised CB to produce 3D reconstructions and brightfield microscopy. ST and CB performed the TEM data acquisition. CB performed the lectin affinity test and immunohistochemistry assays. CB analyzed the data and drafted the first version of the manuscript. TW contributed to all subsequent versions of the manuscript. All authors read, commented and approved the final version of the manuscript.

## Acknowledgments

The authors want to thank the Imaging Unit CIUS, University of Vienna - member of the Vienna Life-Science Instruments (VLSI). JB thanks Andy Sombke (Medical University of Vienna) for providing uranyl acetate and letting us contrast numerous grids in his lab. Furthermore, the authors acknowledge the funding provided by the Faculty of Life Sciences at the University of Vienna.

## Supplementary figures

**Supplementary Figure 1.**
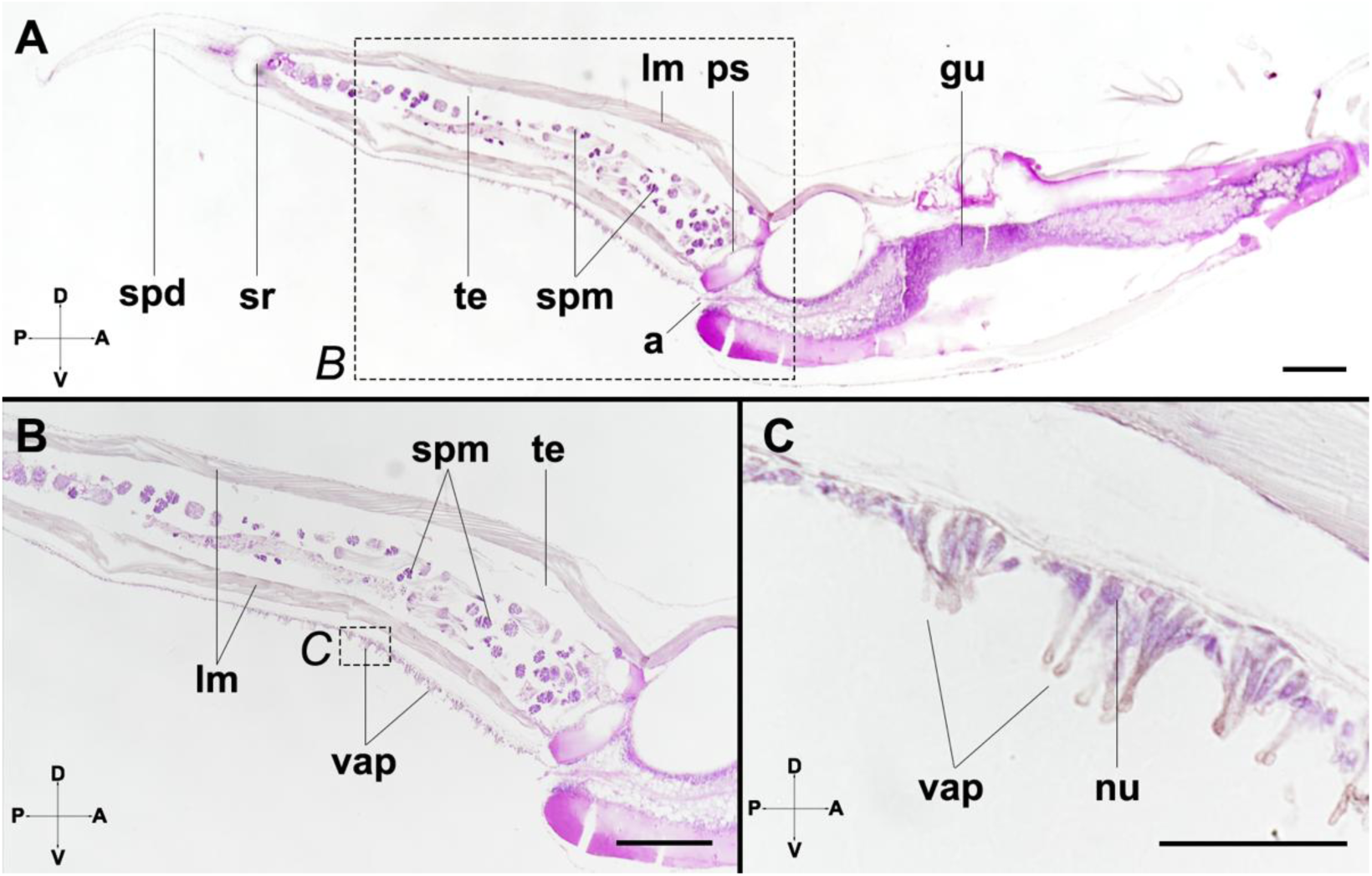
Negative control for the Periodic Acid Shift (PAS) reaction. Amylase digestion before PAS staining the adult of *Spadella cephaloptera*. The staining (violet) indicates PAS^+^ molecules after glycogen digestion and are therefore not included into the discussion. **A.** Sagittal section stained with PAS after amylase digestion. In the tail region, the spermatogonial masses (spm) within the testes (te) exhibit staining. Regarding the trunk, there is strong staining in the gut (gu)**. B.** Detail of the tail region (referenced in A) showing the absence of staining in the testes. **C.** Close-up to the ventral adhesive papillae (vap), where no PAS^+^ labelled molecules are present when amylase treatment is applied. (noted in B) Additional abbreviations: a, anus; lm, longitudinal muscle; nu, nuclei; ps, posterior septum; spd, sperm duct; sr, seminal receptacle. Scale Bars: A, B = 200 µm; C = 50 µm.

**Supplementary Figure 2.**
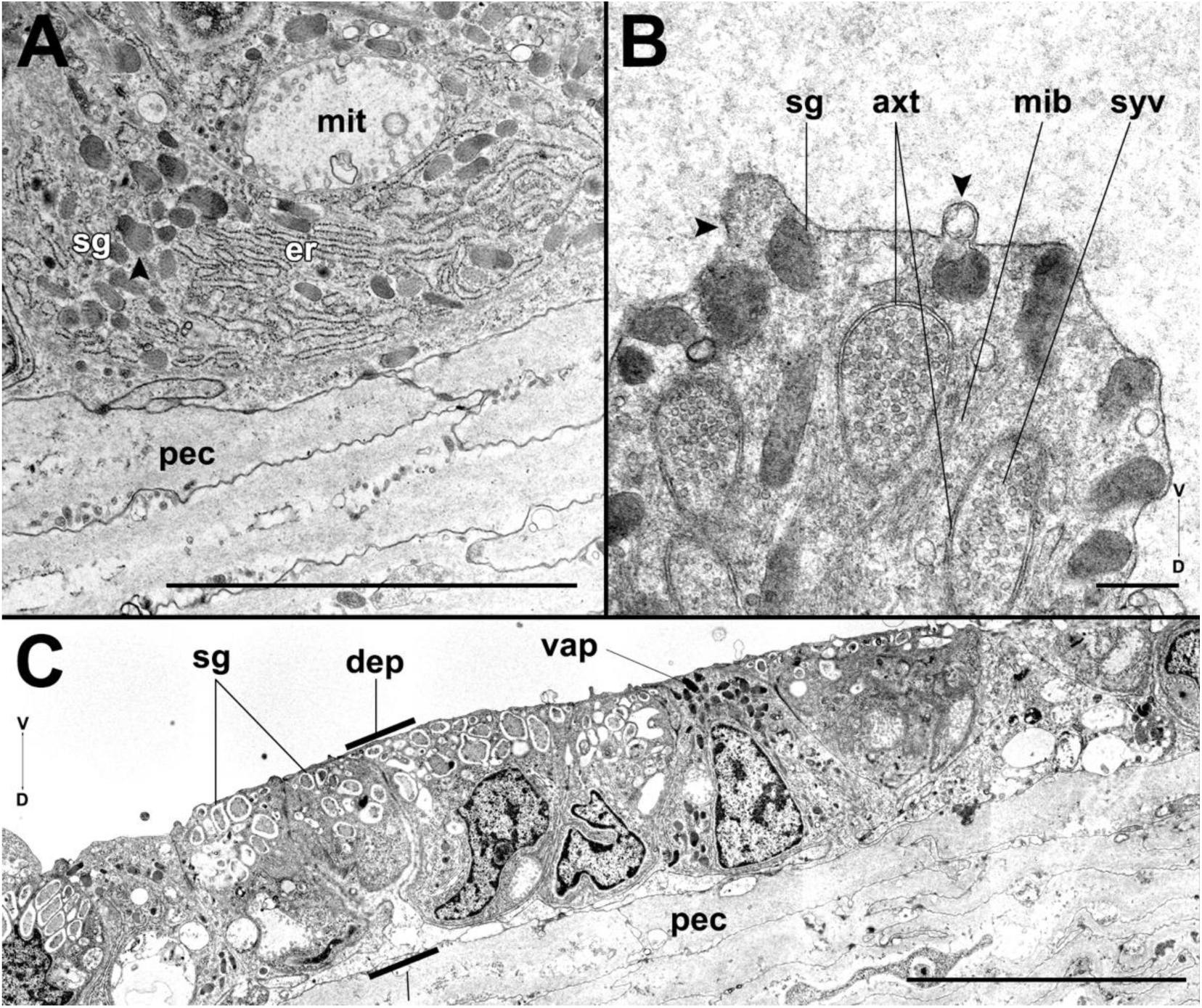
Ultrastructural features of the adhesive cells of *Spadella cephaloptera*. **A.** Detail on the origin of the secretion granules (sg) (arrowhead) in the endoplasmic reticulum (er), localized in the basal region of the adhesive cells. **B.** Indication of exocytosis events of the electron-dense granules in the apical region of the adhesive cells (arrowheads) at 5dph**. C.** Ultrastructure of the distal epidermis (dep), sitting on the proximal epidermis (pec) in the adult chaetognath, where epidermal cells exhibit a high density of secretion granules and surround the ventral adhesive papillae (vap). Additional abbreviations: axt, axonal terminal; mib, microtubules; mit, mitochondria; synaptic vesicles. Scale Bars: A = 5 µm, B = 500 nm, C = 10 µm.

**Supplementary Figure 3.**
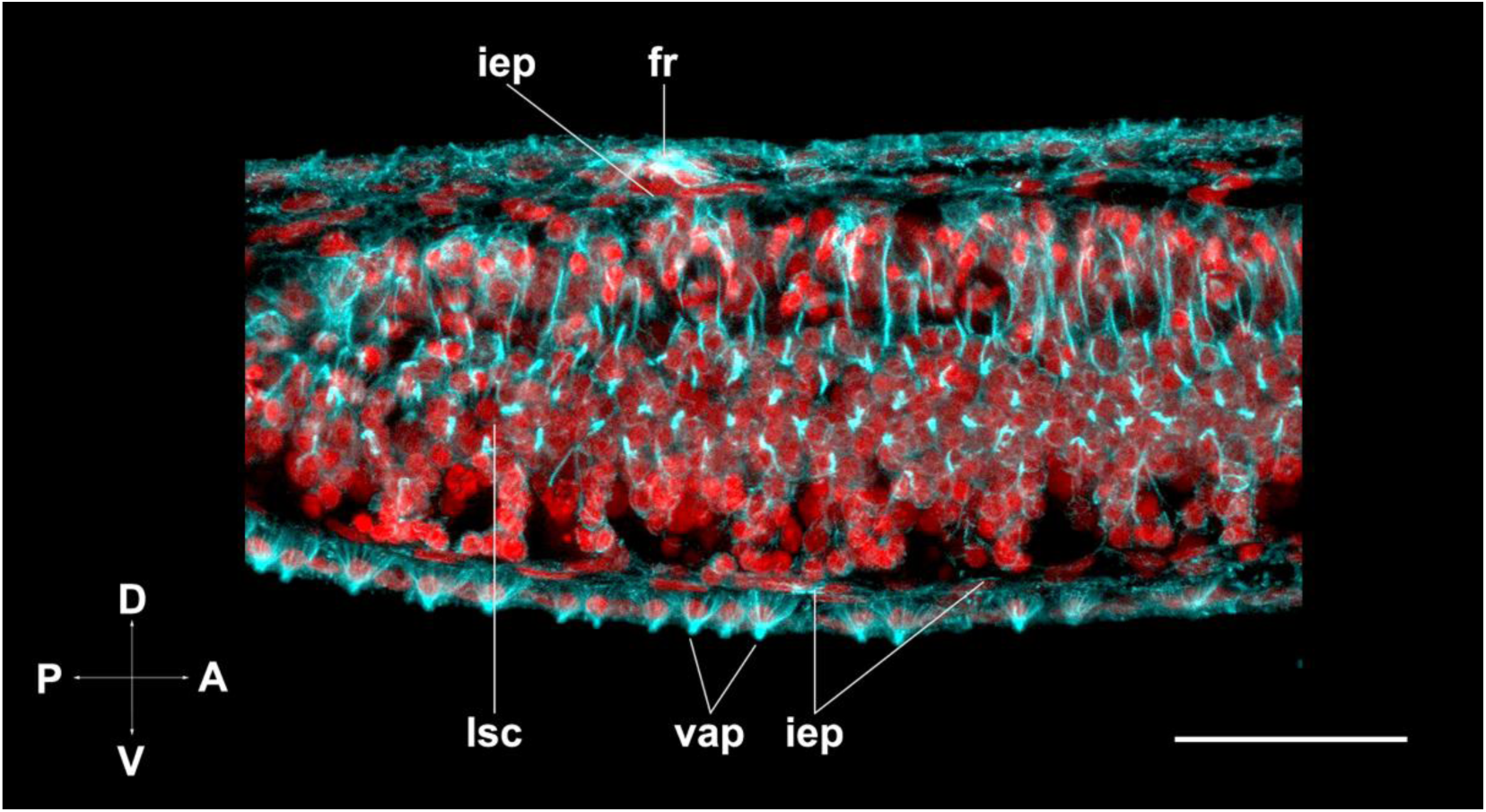
Distribution of acetylated microtubules in the trunk region of *Spadella cephaloptera* at 24 hph. From the lateral somata cell clusters (lsc), acetylated microtubules project into the intraepidermal plexus that connects to multiple cell types in the surface of the epidermis, such as the fence receptors (fr) and the ventral adhesive papillae (vap). Scale Bars: 50 µm.

